# Functional and microstructural plasticity following social and interoceptive mental training

**DOI:** 10.1101/2020.11.11.377895

**Authors:** Sofie L Valk, Philipp Kanske, Bo-yong Park, Seok Jun Hong, Anne Böckler, Fynn-Mathis Trautwein, Boris C. Bernhardt, Tania Singer

## Abstract

The human brain scaffolds social cognitive functions, including Theory of Mind, empathy and compassion, through its functional and microstructural organization. However, it remains unclear how the learning and refinement of these skills may, in turn, shape brain function and structure. Here we studied if different types of social mental training can lead to plastic changes in brain function and microstructure. We studied a group of 332 healthy adults (197 women, 20-55 years) with repeated multimodal neuroimaging and behavioral testing. Our neuroimaging approach capitalized on the quantification of cortical functional gradients and myelin-sensitive T1 relaxometry, two emerging measures of cortical functional organization and microstructure. Longitudinal analysis indicated marked changes in intrinsic cortical function and microstructure, which varied as a function of social training content. In particular, we observed consistent differential change in function and microstructure between attention-mindfulness and socio-cognitive training in regions functionally associated with attention and interoception, including insular and parietal cortices. Conversely, socio-affective and socio-cognitive training resulted in differential microstructural changes in regions classically implicated in interoceptive and emotional processing, including insular and orbitofrontal areas, but did not result in functional reorganization. Notably, longitudinal changes in cortical function and microstructure were predictive of behavioral change in attention, compassion and perspective-taking, suggesting behavioral relevance. In sum, our work provides evidence for functional and microstructural plasticity after the training of social-interoceptive functions, and provides a causal perspective on the neural basis of behavioral adaptation in human adults.

## Introduction

Humans unique social skills enhance cooperation and survival (1, 2). Social capacities can be divided into multiple sub-components (3–5): (i) socio-affective (or emotional-motivational) abilities such as empathy allowing us to share feelings with others, and may give rise to compassion and prosocial motivation (6–8); (ii) socio-cognitive abilities gain access to beliefs and intentions of others [also referred to as Theory of Mind (ToM) or mentalizing (3, 9, 10)]. Finally, interoceptive abilities, attention, and action observation serve as important auxiliary functions of social aptitudes, contributing to self-other distinction and awareness (11–13). Together, these capacities combine externally and internally oriented cognitive and affective processes and reflect both focused and ongoing thought processes (14–18). With increasing progress in task-based functional neuroimaging, we start to have an increasingly precise understanding of brain networks associated with the different functional processes underlying social cognition. For example, tasks probing socio-emotional functioning and empathy consistently elicit functional activations in anterior insula, supramarginal gyrus, and midcingulate cortex (3, 19, 20), while emotional-motivational processes, such as compassion, implicate insular and orbitofrontal areas (21, 22). On the other hand, tasks involving socio-cognitive functioning generally activate regions of the human default mode network (DMN), such as medial frontal cortex, temporo-parietal junction, and superior temporal sulcus (4, 10, 23). Finally, attentional tasks activate inferior parietal and lateral frontal and anterior insular cortices (24–26) and interoceptive awareness is linked to anterior insula and cingulate regions (12, 13, 27, 28). Together, these findings suggest a potentially dissociable neural basis of different social abilities in the human brain.

Despite the progress in the mapping of the functional topography of networks mediating social and interoceptive abilities, there remains an overall gap in the understanding of the specific microarchitecture of these networks (29). In general, however, prior research has shown that cortical function and microstructure follow parallel spatial patterns, notably sensory-fugal gradients that may support the differentiation of sensory and motor function from higher order functional processes, such as social cognition (30–34). Put differentially, a sensory-transmodal gradient approach situates higher social and interoceptive functions its transmodal anchors, encompassing both heteromodal regions (such as the prefrontal cortex, posterior parietal cortex, lateral temporal cortex, and posterior parahippocampal regions) as well as paralimbic cortices (including orbitofrontal, insular, temporo-polar, cingulate, and parahippocampal regions) (35). Distant from sensory systems, these transmodal systems take on functions that are only loosely constrained by the immediate environment (36), allowing internal representations to contribute to (more abstract, and social) cognition and emotion (32, 33, 36–43), thereby enhancing behavioral flexibility (38, 44). However, despite the presumed link between cortical microstructure and function, how changes in social behavior impact brain function and microstructure it is not known to date.

Longitudinal investigations can reveal causal links between behavioral skills and brain organization, for example via targeted mental training studies. In the context of mental skills, a range of prior studies indicated that mental training may alter brain function and gross brain morphology (45–49), but findings do not yet point to a consistent pattern. For example, a randomized controlled trial showed little effect on brain morphology of 8 weeks of mindfulness-based training in healthy adults (50). In general, however, sample sizes have been relatively modest and training intervals short. Moreover, few studies have compared different practices or focussed on different social skills specifically, though it is likely different types of mental training have unique effects on brain and behavior (51–53). In a previous study realized in the context of the *ReSource* project (54), our group could show differentiable change in MRI-derived cortical thickness, in support of macrostructural plasticity of the adult brain following the training of different social and interoceptive skills (55). In brief, the ReSource project involved a targeted training of attention-mindfulness (*Presence* module), followed by socio-affective (*Affect* module) and socio-cognitive/ToM training (*Perspective* module) over the course of nine months. Whereas *Presence* aimed at initially stabilizing the mind and nurturing introspective abilities, the *Affect* and *Perspective* modules focussed on nurturing social skills such as empathy, compassion, and perspective taking on self and others.

Here, we leverage the *ReSource* study dataset to assess whether the targeted training of attention-interoception, socio-affective, and socio-cognitive skills can lead to reorganization of (i) intrinsic function (as indexed by resting-state fMRI connectivity gradient analysis), and (ii) cortical microstructure (as indexed by quantitative T1 relaxometry, probed along the direction of cortical columns (56–58)). Such results would be in line with prior observations suggesting coupled change in brain structure and function (59, 60), and would help to gain insights into the association between social skills and brain organization. In particular, we carried out longitudinal analyses of subjects-specific measures of functional integration and segregation and evaluated whether these changes corresponded to longitudinal change in intracortical microstructure. We hypothesized that changes in functional activity, as captured by intrinsic functional brain organization, relate to changes in intracortical microstructure. In addition to assessing the reorganization of cortical microstructure and function, we measured associations to behavioural change in attention, compassion, and ToM markers using machine learning with cross-validation to evaluate behavioural relevance of the observed changes at the individual level.

## Results

### Embedding of socio-affective and -cognitive functions in cortical brain organization (Fig. 1)

**Figure 1.**
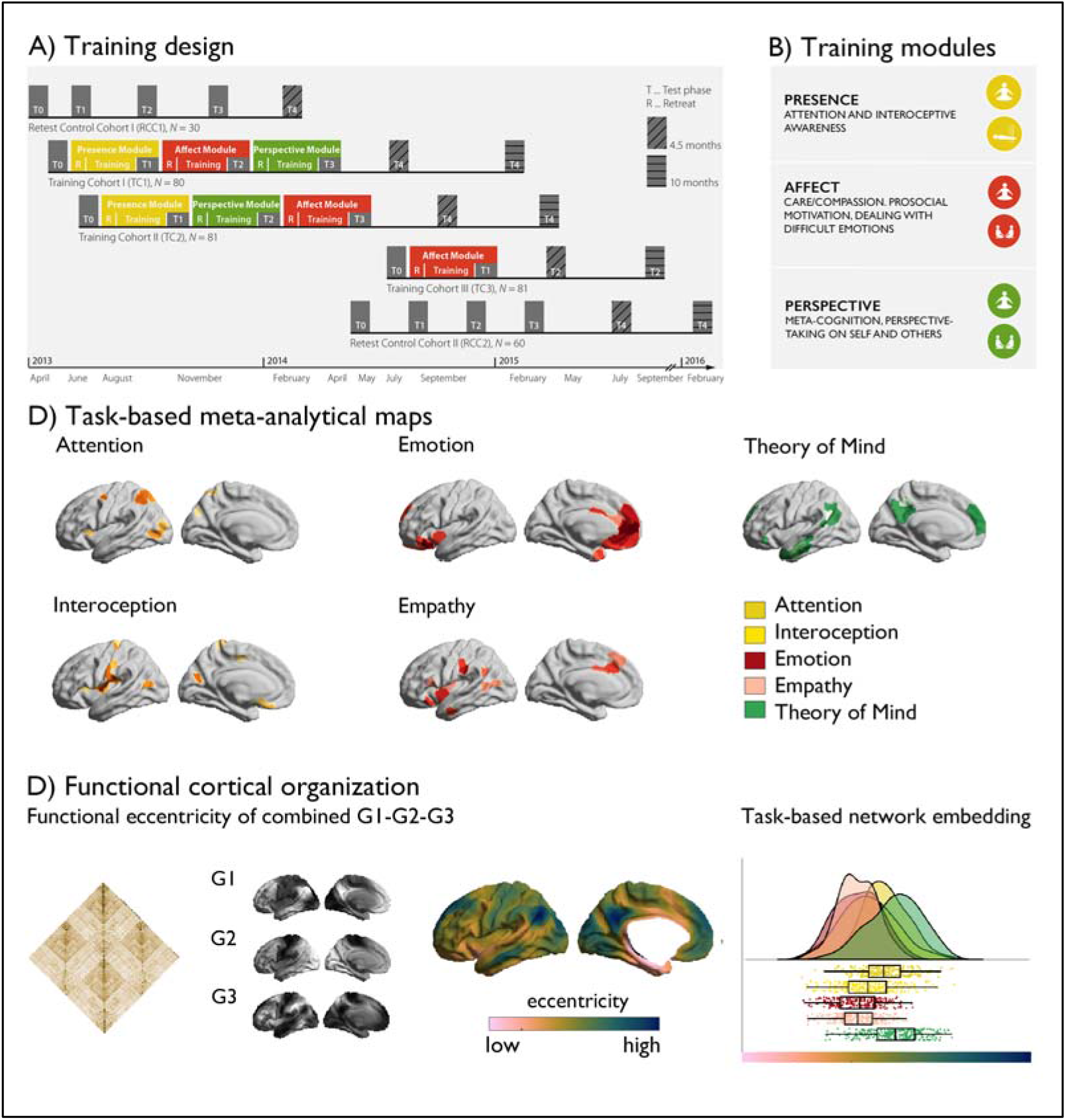
**A)** Training design of the ReSource study; **B**) Training modules; **C**) Task-based meta-analytical maps, and a legend of the color-coding of the maps; **D**) Functional cortical organization, from left to right: functional connectivity matrix, gradient 1 – 3, eccentricity metric and task-based network embedding.

We analyzed resting-state functional MRI (fMRI) measures, myelin-sensitive quantitative T1 (qT1) relaxometry, and behavioral data from 332 adults studied in the *ReSource Project* (54). The preregistered trial (https://clinicaltrials.gov/ct2/show/NCT01833104) involved three 3-month long training modules: (i) *Presence*, targeting interoception and attention, (ii) *Affect*, targeting empathy and emotion, and (iii) *Perspective*, targeting ToM.

For *a-priori* functional localization, we selected meta-analytical functional networks mapping these functions using NeuroSynth (61), (**Figure 1**). Cortex-wide functional organization was then assessed after identifying spatial gradients of resting-state functional connectivity (36, 62–65), derived using open access techniques (62). Gradients of each individual were Procrustes aligned to the mean functional connectome based on the human connectome project sample (30, 66), and we calculated region-wise distances to the center of a coordinate system formed by the first three gradients G1, G2, and G3 for each individual [based on the Schaefer 400 parcellation (67)]. Such a gradient eccentricity measures captures intrinsic functional integration (low eccentricity) vs segregation (high eccentricity) in a single scalar value (68). Highest segregation was observed in visual and sensory-motor networks, while ventral attention and limbic networks were closest to the center of the space. Notably, the *a-priori* networks showed a unique embedding in gradient space (F(5,394) 8.727, p<0.001), with Affect-associated networks being most integrated while and *Perspective*-networks were most segregated.

### Mental training-specific change in functional eccentricity

We then tracked training-related longitudinal changes in the functional gradients G1, G2, and G3 following the different *ReSource* modules. In the *Resource* study, participants were randomly assigned to two training cohorts (TC1, N=80; TC2, N=81), which each underwent a 9-month training consisting of three sequential modules (*i.e*., Presence, Perspective, and Affect) and with weekly group sessions and daily exercises, completed via cell-phone and internet platforms (**Figure 1, Table 1–3**, *Materials and Methods* and *Supplementary Materials* for details). TC1 and TC2 underwent the latter two modules in different order (TC1: Affect→Perspective; TC2 Perspective→Affect) to serve as active control groups for each other (**Figure 1A**). Additionally, a matched test-retest control cohort did not undergo any training (RCC, N=90), but was followed with the same neuroimaging and behavioral measures as TC1 and TC2. All participants were measured at the end of each three-month module (T_1_, T_2_, T_3_) using 3T MRI and behavioral measures that were identical to the baseline (T_0_) measures. There was furthermore an active control group (TC3; N=81), which completed three months of *Affect* training only.

**Table 1.** Participant inclusion in resting-state analysis and quantitative T1 analysis.

| Recruited (N, mean age, % female) | T<sub>0</sub> (N) | T<sub>1</sub> (N) | T<sub>2</sub> (N) | T<sub>3</sub> (N) |
| --- | --- | --- | --- | --- |
| Total (N = 332) | 268 | 259 | 182 | 184 |
| TC1 (N = 80; 41.3; 58.8) | 69 | 65 | 57 | 55 |
| TC2 (N = 81; 41.2; 59.3) | 67 | 59 | 61 | 64 |
| RCC (N = 90; 40.0; 58.9) | 65 | 68 | 64 | 65 |
| TC3 (N = 81; 40.4; 60.5) | 67 | 67 |  |  |

**Table 2.**
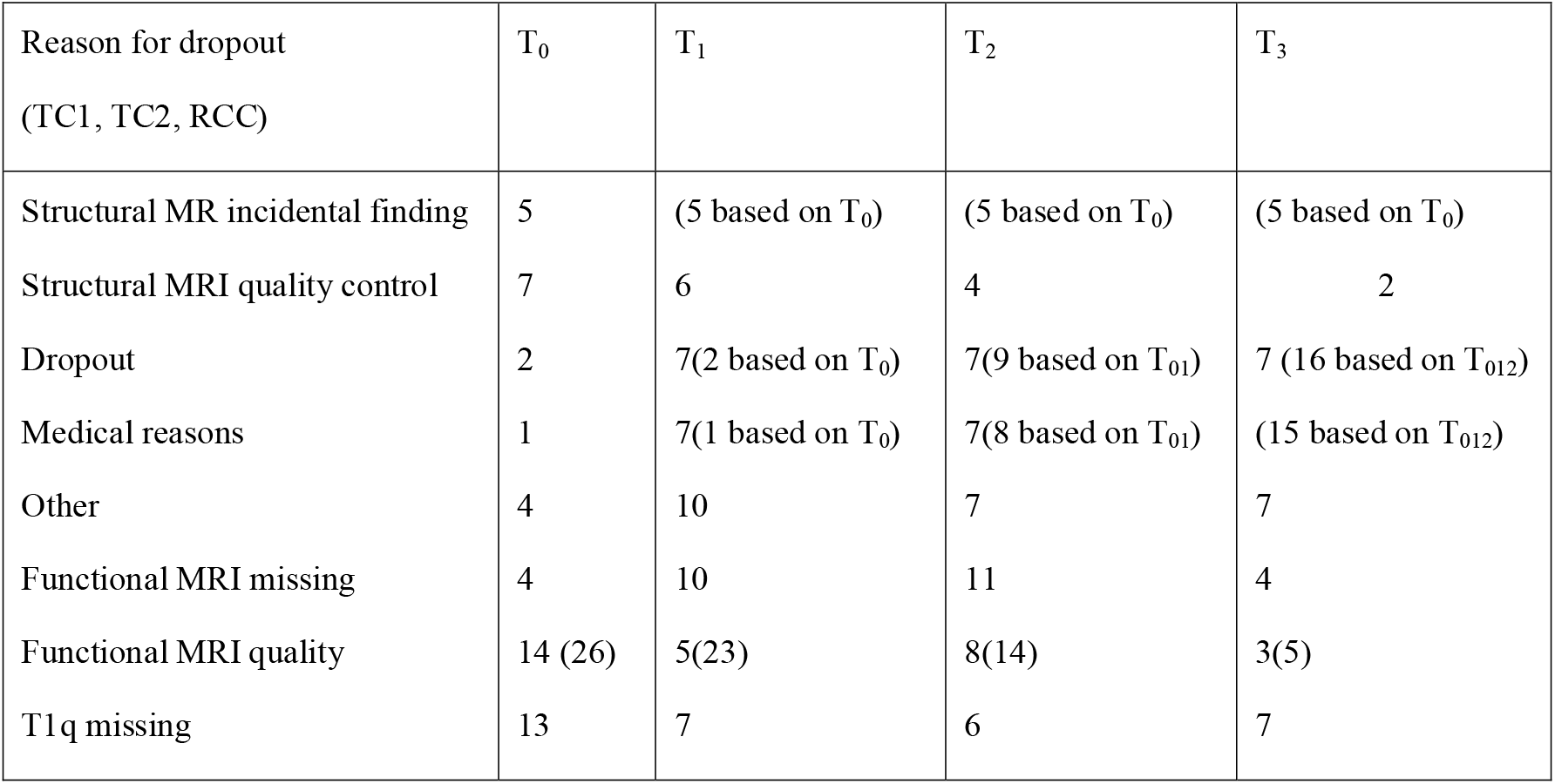
Reason for missing data across the study duration. *MR incidental findings* are based on T_0_ radiological evaluations; participants who did not survive *MRI quality control* refers to movement and/or artefacts in the T1-weighted MRI; dropout details can be found in (54); *no MRT*: due to illness / scheduling issues / discomfort in scanner; *other*: non-disclosed; *functional MRI missing*: no complete functional MRI; *functional MRI quality*: >0.3mm movement (low quality in volume + surface)

**Table 3.**
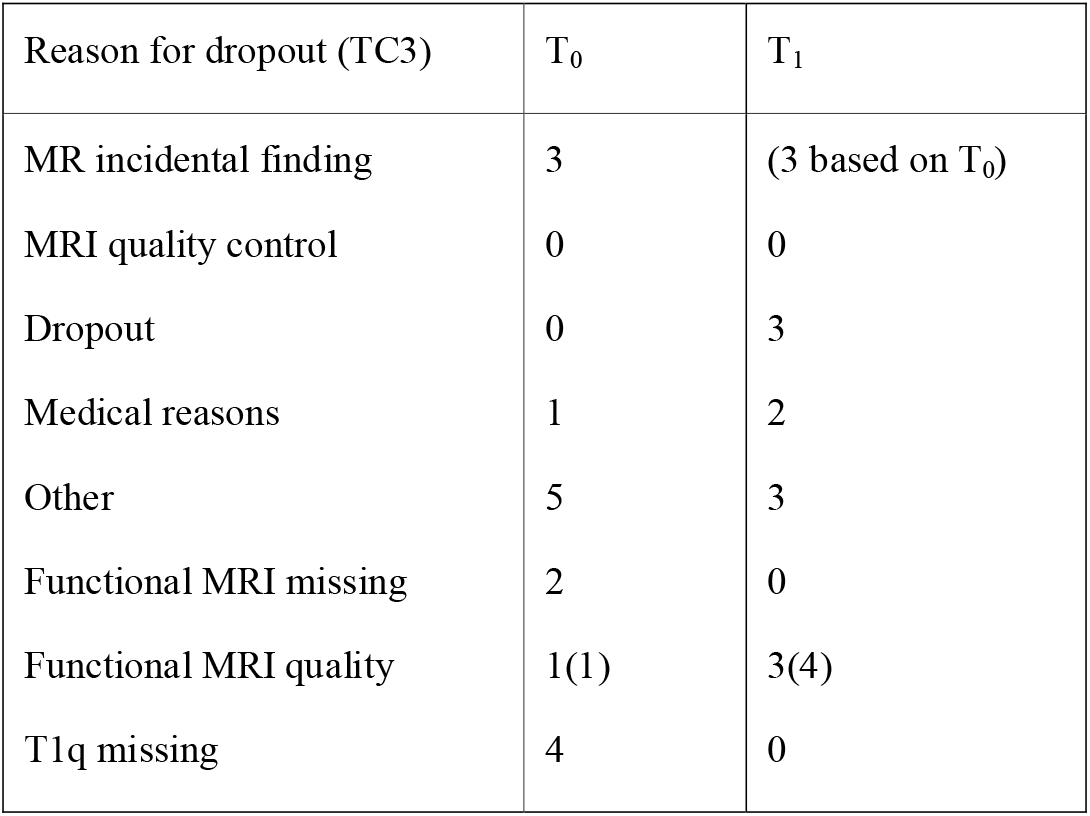
Reason for missing data across the study duration. *MR incidental findings* are based on T_0_ radiological evaluations; participants who did not survive *MRI quality control* refers to movement and/or artefacts in the T1-weighted MRI; dropout details can be found in (54); *no MRT*: due to illness / scheduling issues / discomfort in scanner; *other*: non-disclosed.

| Reason for dropout (TC3) | T<sub>0</sub> | T<sub>1</sub> |
| --- | --- | --- |
| MR incidental finding | 3 | (3 based on T<sub>0</sub>) |
| MRI quality control | 0 | 0 |
| Dropout | 0 | 3 |
| Medical reasons | 1 | 2 |
| Other | 5 | 3 |
| Functional MRI missing | 2 | 0 |
| Functional MRI quality | 1(1) | 3(4) |
| T1q missing | 4 | 0 |

We evaluated how cortical functional gradients would change following mental training using mixed-effects models (69). Excluding participants with missing functional or structural data, or excessive movement, the sample included 109 individuals for *Presence*, 104 individuals for *Affect*, 96 individuals for *Perspective*, 168 *retest controls* and 60 *active controls* (*Affect*) with functional and structural change scores. At the whole-cortex level, comparing all training groups and Modules, we observed marked gradient eccentricity changes following *Presence* and *Perspective* (**Figure 2, Supplementary Table 1-4**). *Presence* training resulted in increased eccentricity of bilateral temporal and right superior parietal areas (FDRq<0.05), indicative of increased segregation. *Perspective* training resulted in decreased eccentricity of right temporal regions, together with left insular cortices (FDRq<0.05). We observed no eccentricity change following *Affect* training. *Post-hoc* analysis indicated changes between *Presence* and *Perspective* were underlying eccentricity change were most marked in G2 (t= - 4.647, p<0.001, d=-0.403), dissociating sensory-motor from visual systems, but not G1 (t= - 1.495, p>0.05, d=-0.130) or G3 (t= −0.493, p>0.05, d=-0.043) gradient. Focussing on a-priori networks, in particular attention (t=2.842, p=0.005, d=0.247) and interoception (t=2.765, p=0.006, d=0.240) networks showed alterations in *Presence-vs-Perspective*, **Table 4, Figure 2 and Supplementary Figure 1**. Though effects varied, they were also observed after GSR control, in TC1 and TC2, and versus RCC (**Supplementary Table 5-9**). Evaluating gradient-specific alterations per a-priori network we observed a link between *Presence* versus *Affect* in the empathy-network along G2 (t=3.215, p<0.002) **(Supplementary Table 10-12, Supplementary Figure 2-4)**. Findings were robust when controlling for previously reported cortical thickness change (55), **Supplementary Table 13**. We did not find evidence for overall effects of training on functional eccentricity relative to RCC (**Supplementary Table 14**).

**Table 4.** Module-specific changes in eccentricity per functional networks. * signifies FDR corrected differences.

|  | Presence vs Perspective | Presence vs Affect | Perspective vs Affect |
| --- | --- | --- | --- |
| Attention | $t = 2.842$, $p = 0.005^*$, $d = 0.247$ | $t = 1.458$, $p > 0.05$, $d = 0.127$ | $t = -1.692$, $p > 0.05$, $d = -0.147$ |
| Interoception | $t = 2.765$, $p = 0.006^*$, $d = 0.240$ | $t = 1.043$, $p > 0.05$, $d = 0.091$ | $t = -2.008$, $p = 0.045$, $d = -0.174$ |
| Emotion | $t = 0.387$, $p > 0.05$, $d = 0.035$ | $t = -0.135$, $p > 0.05$, $d = -0.011$ | $t = -0.552$, $p > 0.05$, $d = -0.048$ |
| Empathy | $t = 2.218$, $p = 0.027$, $d = 0.193$ | $t = 0.879$, $p > 0.05$, $d = 0.076$ | $t = -1.569$, $p > 0.05$, $d = -0.136$ |
| Theory of Mind | $t = 1.721$, $p > 0.05$, $d = 0.149$ | $t = 1.324$, $p > 0.05$, $d = 0.115$ | $t = -0.601$, $p > 0.05$, $d = -0.052$ |

**Figure 2.**
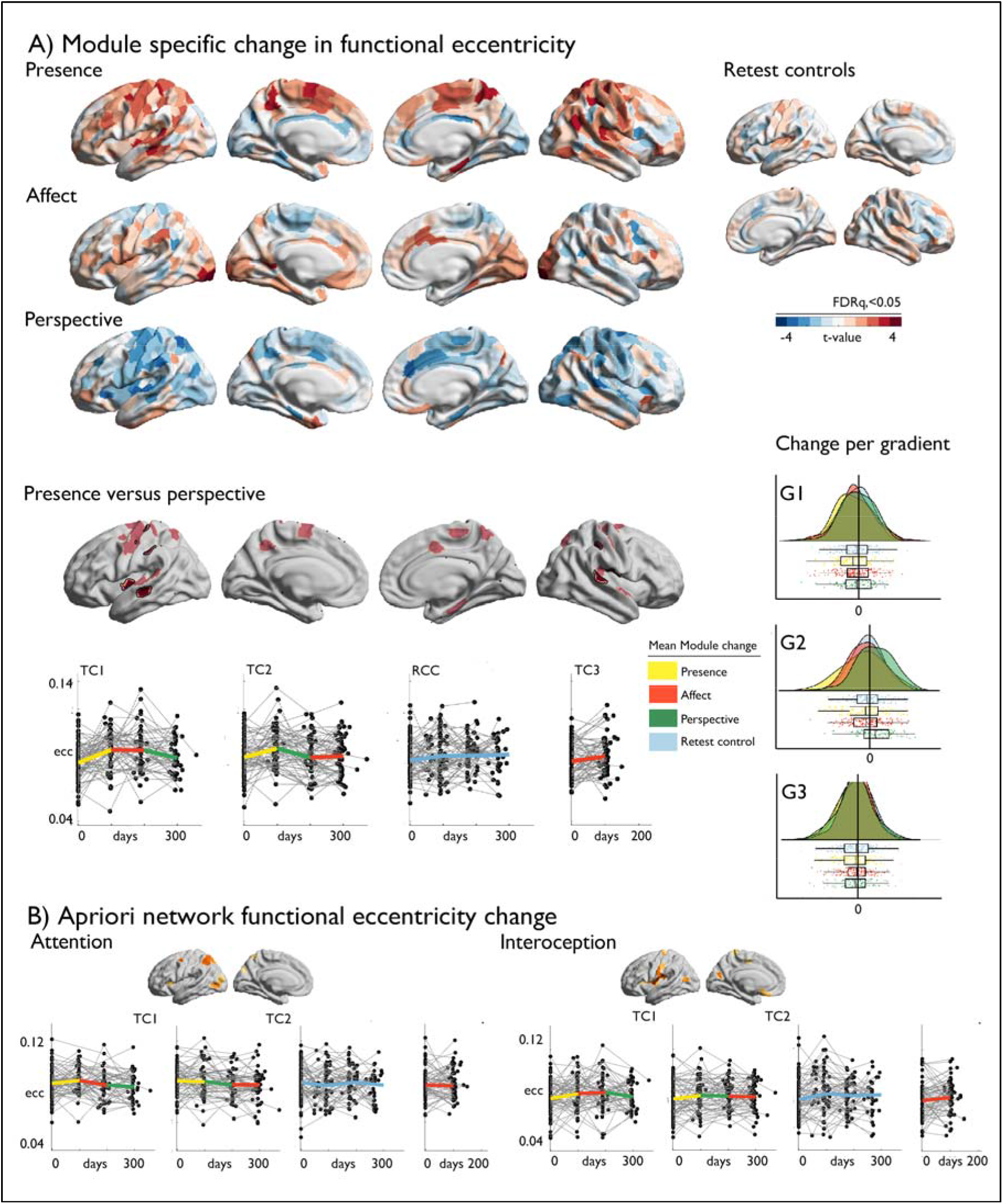
Training-induced changes in cortical functional organization. **A**) *T*-maps of module-specific changes in functional eccentricity; **B**) Module-specific change in functional eccentricity, p<0.01, FDRq<0.05 outlined in black, *below*: alterations of eccentricity in the FDRq<0.05 regions, *right*: mean changes in FDRq<0.05 eccentricity regions as a function of G1-G2-G3; **C**) A-priori network functional eccentricity change in networks that showed Module specific change.

### Dissociable microstructural alterations following mental training

Having shown alterations in functional gradients following social mental training, we aimed to evaluate whether there are corresponding changes in cortical microstructure. To do so, we sampled qT1 relaxometry values across 12 equidistant intracortical surfaces between the pial and the white matter (32), **Figure 3**. Regions with low mean qT1 were located in sensorymotor and visual regions, regions known to have a high myelin content (70, 71). On the other hand, regions with high mean qT1 and thus low myelin content were located in transmodal areas, as previously shown (57). We then examined how intra-cortical microstructural organization mirrored observed changes in functional eccentricity in clusters showing differential change during *Presence* vs *Perspective*. We observed a correspondence (FDRq<0.05) between functional eccentricity and upper layer microstructural compartments (1st: t= 3.167 p=0.002, d=0.275; 2nd compartment: t= 2.911, p=0.004, d=0.253) in regions showing differences in eccentricity between *Presence* and *Perspective*. Following, assessing training module-specific effects in microstructure in *a-priori* task-based functional networks through comparing all training groups and Modules, we found all but the emotion network to show increases in qT1 of *Presence* versus *Affect* and *Perspective* in the upper compartments, extending to deeper compartments when comparing *Presence* and *Affect* (FDRq<0.05). Conversely, in deeper compartments, near the GM/WM boundary, we observed decreases of *Affect* relative to *Perspective* in interoception and emotion-related networks (**Supplementary Table 15-21, Supplementary Figure 5**). Findings were largely consistent across the different training cohorts, yet weak relative to retest controls (**Supplementary Table 22-35, Supplementary Figure 6 and 7**). As for the functional change, findings were also observed when controlling for cortical thickness (**Supplementary Table 36-38**), indicating that microstructural change goes above and beyond previously reported morphological change (55). Overall, *ReSource* training led to decreased qT1 values, i.e. increased myelination, in both TC1 and TC2 relative to RCC over the nine months training time, in all functional networks in particular in deeper layer microstructure, whereas RCC showed subtle increases of qT1, suggesting decreased myelination (**Supplementary Table 39, Supplementary Figure 8**). Exploring correspondence between functional and microstructural change within Modules, rather than by contrasting Modules, we observed a spatial correlation between functional change in eccentricity and G2 in upper and middle compartment microstructure in *Presence* and overall correspondence with G3 changes, correcting for spatial autocorrelation (p_spin_<0.05), whereas microstructural alterations in mid- and deeper compartments showed correspondence to eccentricity and G2 in *Affect* (p_spin_<0.05).

**Figure 3.**
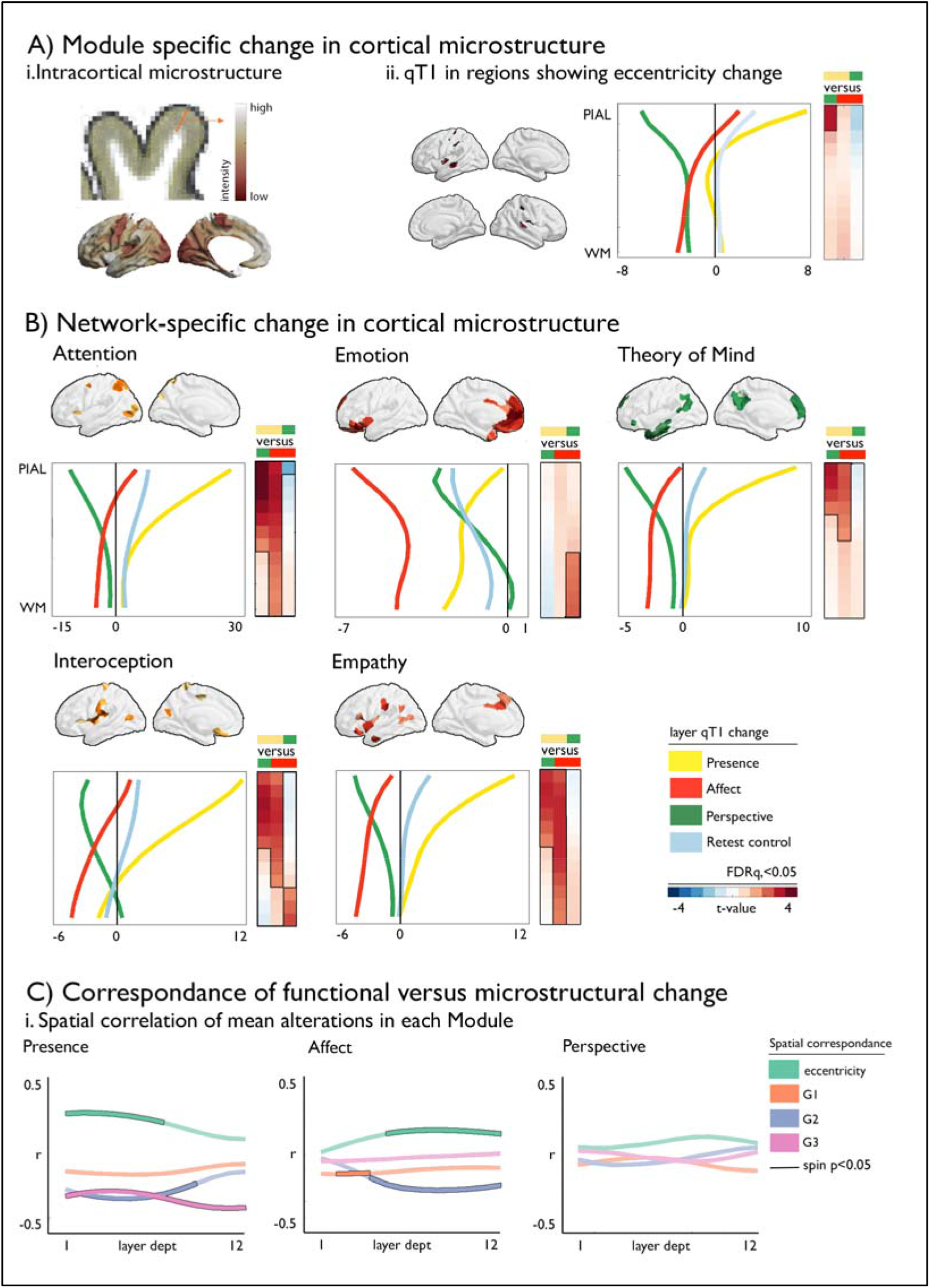
Dissociable microstructural alterations following mental training_. **A**). Module specific changes in cortical microstructure; i. probing dept-specific cortical microstructure and dept-specific mean microstructural differences across functional networks; ii. qT1 in regions showing eccentricity change; **B**) Network-specific change in cortical microstructure as a function of dept, mean change per module is depicted as a spaghetti plot, p_FDR_<0.05 have black outline; **C**) Correspondence of functional versus microstructural change; i. Spatial correlation of mean alterations in each Module, black outline indicates p_spin_<0.05.

### Training module-specific prediction

Last, we evaluated whether alterations in cortical microstructure and function following mental training could predict behavioral changes following the *Resource* training. Previous work has indicated Module specific behavioral changes in attention, compassion, and perspective-taking, as measure using a cued-flanker (attention) and the EmpaTom task (compassion and perspective-taking)(72). Supervised learning (lasso regression, 5-fold cross validation, 100 repetitions) with sequential feature selection (7 components, 20% of features) was utilized to predict behavioral change from the average functional gradient eccentricity, and G1-G3, as well as microstructure in superficial (1:4), middle (5:8) and deep (9:12) compartments in the five functional networks, resulting in 35 features to select from (**Figure 4**). Attention predictions (N=85, TC1 and TC2) were most marked in attention network in microstructure at superficial depts and eccentricity (MAE (mean±SD): −0.037±0.003, out of sample r (mean±SD): 0.325±0.308). Conversely, compassion (N=100, TC1 and TC2) was predicted primarily through structural and functional reorganization of attention, interoception and emotion networks (MAE: −0.412±0.029, out of sample r: 0.284±0.279). Last, Theory of Mind (N=93, TC1 and TC2) predictions (MAE: −0.081±0.006 out-of-sample r: 0.301±0.281) were most likely to occur in attention networks along G3 and microstructure of upper and middle compartments, as well as emotion related networks along G3.

**Figure 4.**
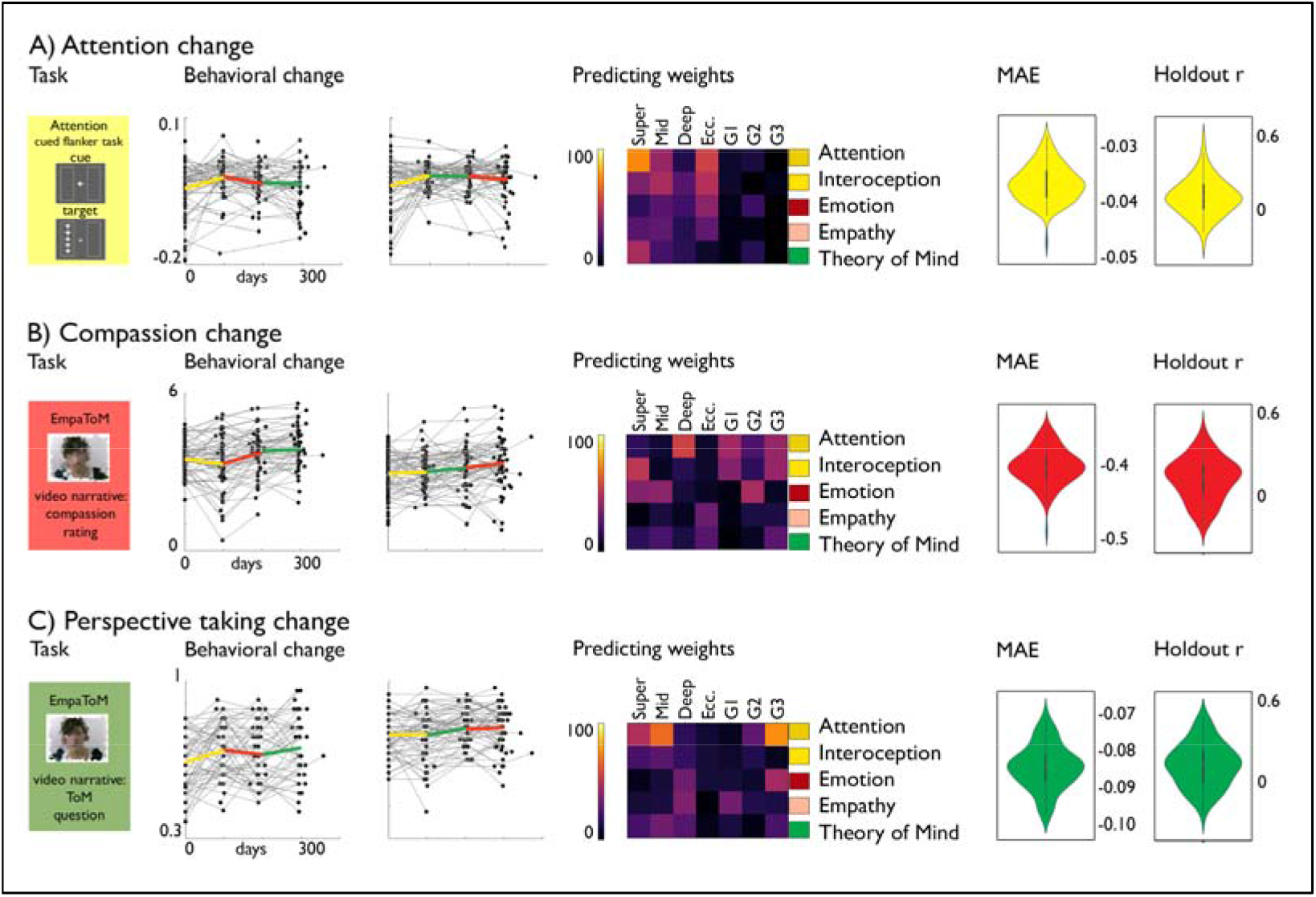
**A**). Attention change; from *left* to *right*: attention task, behavioral change, predicting weights, MAE and holdout r distribution; **B**). Compassion change; from *left* to *right*: compassion task, behavioral change, predicting weights, MAE and holdout r distribution; **C**). Perspective-taking change; from *left* to *right*: perspective-taking task, behavioral change, predicting weights, MAE and holdout r distribution.

## Discussion

Our main goal was to deepen our understanding of the functional and structural plasticity of the human cerebral cortex by focusing on how cultivating cognitive and social skills such as attention, empathy, compassion and perspective taking through mental training can systematically alter both functional and microstructural cortical organization. To achieve this goal, we evaluated longitudinal changes in MRI-derived cortical functional gradients and qT1 profiles as well as their interrelationship in the context of the 9-month *ReSource* study (54). Our study provides evidence of functional and microstructural plasticity that varied as a function of distinct social mental trainings, and that can predict training-related behavioural change.

Our results demonstrate change in intrinsic functional architecture following the training of social and attentional functions. Particularly, our findings show increased functional segregation (as indicated by increases in functional gradient eccentricity) of regions in attention and interoceptive networks, including parietal and posterior insular regions, following *Presence* training, indicating that these networks functionally differentiated from the rest of the cortex. Conversely, *Perspective* training resulted in an increased functional integration of these regions with other brain networks. These observations were mirrored by alterations in cortical microstructure between both training Modules, in particular in upper layers of the cortex. Moreover, we found microstructural change in deeper compartments in regions associated with interoception and emotion processing, including insular and orbito-frontal regions, when contrasting *Perspective* and *Affect* Modules. Last, we could predict training-specific increases in attention, compassion and ToM scores based on co-occurring changes in large-scale functional and dept-varying microstructural organization, suggesting behavioral relevance of functional and microstructural alterations following mental training. In sum, our study showed that the concerted training of social skills results in dissociable changes in the functional connectome with corresponding alterations in cortical microstructure. Notably, our work indicates that the human brain changes as a function of mental training content, not only as a function of spatial distribution of regions along the cortical mantle, but also as a function of its functional organization axes and cortical depth. This provides novel and longitudinal evidence of the interrelationship of functional organization in intracortical microstructure in the context of human cognition.

Analyzing meta-analytical fMRI networks involved in social and attentional processes at baseline, we could show that networks associated with social processing were differentially positioned in a coordinate system spanned by the first three functional gradients. Echoing a mounting literature (36, 65, 73, 74), the principal gradient ran from unimodal to transmodal systems with as apex the DMN. This axis aligns with classic notions of cortical hierarchy (38, 75), axes of microstructural differentiation (32, 33, 57) and cortical evolution, with heteromodal regions undergoing recent expansions in the human lineage (34, 76–78). Conversely, the second gradient dissociates visual and sensory systems and the tertiary gradient dissociates the task-positive, attention and control, networks from rest of the brain (36). Unlike the DMN, the task-positive network, including frontal and parietal regions, engages preferentially in externally oriented tasks (79–81). Thus, this axis may describe a differentiation between DMN-related socio-episodic memory processing from task-focused processing associated with the multiple demand network (16, 82). Together the three gradients describe an organization of integrated and segregated functional processing, with primary and default regions showing functional segregation and saliency network functional integration (68, 83, 84). Indeed, along its axes, we found functional networks associated with ToM and attention to be most segregated, indicating that their functional architecture is relatively distinct from other transmodal networks. Conversely, emotion-related, and to a, lesser extend empathy and interoceptive processing, functional networks were found to be relatively integrated with all other networks. This positioning may be reflective of a regulative role of functional activity of task-positive and -negative networks (85). As such, we could show that the meta-analytical functional activations associated with social cognition each were placed at unique locations along cardinal axes of functional organization, showing varying levels of functional integration and segregation.

Studying functional network plasticity following *ReSource* training, we found that regions associated with attention and interoception showed functional integration following *Perspective* training, whereas they segregated following *Presence*. In the context of social cognition, integrated states might reflect active thought processes such as those required when engaging in ToM, while more segregated states may reflect automation and domain-specific function (86). Indeed, attention-based mindfulness, as cultivated during *Presence*, may reduce habitual thought patterns and enhance momentary awareness (87) – possibly captured by functional network segregation. Moreover, our observations in the domain of social skill training align well with a previous observation that working memory task performance is associated with decreased modularity (increased integration), whereas automation following working memory training was associated with increased modularity (segregation) of multiple demand network and DMN (86). Second, 8-week simultaneous training of compassion, mindfulness and perspective taking has been reported to result in reduced intra-network connectivity in the DMN, VAN, and somatomotor networks, reflecting integration (88). More generally, our findings may be in line with the Global Workspace Theory, which poses that automated tasks, such as interoception and awareness, can be performed within segregated clusters of regions, whereas those that are challenging, for example perspective-taking, require integration (89). Of note, we did not observe changes in eccentricity following the *Affect* training module, relative to the other modules. Rather, *Affect* seemed to stabilize changes in eccentricity observed following the other trainings. It is possible that the lack of change in eccentricity following socio-affective training reflects a coordinating role of socioaffect relative to alterations associated with attention-mindfulness and socio-cognition. Such an interpretation aligns with theories of emotional allostasis, suggesting that affective processing may balance integration and segregation of brain function to regulate resources dynamically (14, 15). Notably, though the current work focused on cortical networks, it is likely also subcortical regions contribute to the functional organization of the social brain (90, 91). Follow-up work that studies plasticity of sub-cortical function and structure in the context of the social brain may provide additional system-level insights.

To further evaluate potential neurobiological substrates of cortical functional reorganization, we leveraged an intra-cortical microstructure using equidistant probes perpendicular to the cortical mantle (92, 93). We were particularly interested to study changes in cortical myeloarchitecture, given recent reports of experience-dependent myelin plasticity (59, 60) as well as our previous finding of macrostructural change in the same cohort (55). Evaluating the functional eccentricity alterations in attention and interoception networks observed between *Presence* and *Perspective*, we found that these changes could be particularly recapitulated by microstructural alterations in superficial compartments of the cortex. Whereas socio-cognitive training resulted in increased myelination proxy (reflected by decreases in qT1) values, attention-based mindfulness training resulted in decreased myelination of these regions. Similar patterns were observed in empathy and Theory of Mind networks. Moreover, contrasting attention-mindfulness and social affective training, we observed similar patterns of changes in microstructure in superficial compartments, which extended to middle compartments for attention, interception and empathy-related meta-analytical functional networks. Conversely, we found differential alterations of qT1 comparing socio-cognitive with affective training in deep compartments of interoceptive and emotion-related functional networks, where socio-affective training resulted in relative increase in myelin as compared to socio-cognitive training. Various studies have reported training-induced changes in microstructure in humans, often measured using diffusion MRI derivative measures, such as fractional anisotropy (94–96). Moreover, work in mice has shown that social isolation during development resulted in changes in mPFC myelination, correlating with working memory and social behaviors (97). The observed alterations in qT1 in our study may reflect changes in oligodendrites as well as supporting glial cells (98, 99). Indeed, myelin-plasticity has been suggested to be a key biological marker of learning and memory in adulthood (60, 100). Due to the ‘sub-optimal’ myelination of association regions conduction velocities may still be adjusted dynamically in adulthood, as a function of changing environmental demands(59). Interestingly, alterations persisted above and beyond cortical thickness change alone (55). As structural plasticity includes both synaptic and myelin-related processes, further study on the plasticity of multiple microstructural markers *in vivo* may help to disentangle the different biological processes that underlie adaptive cognition in adulthood.

Co-alterations of function and structure following different types of mental training modules were further evidenced by module specific analyses, including behavioral predictions. Together, they suggest co-alterations of function and microstructure to occur in a manifold fashion, with differential change occurring both in spatially divergent functional networks and as a function of different functional axes and cortical depts. By integrating functional organization with intra-cortical microstructure in the context of a longitudinal study involving social and cognitive mental training in adults, the current study integrates perspectives on structure and functional organization of the human cortex and its relation to cognition. The layered structure of the human cortex is a key element supporting its function, leading up to different functional organizational dimensions, supporting cognition (16, 34, 82). In particular, the layered structure of the cortex enables feedforward and feedback information transfer between regions, supporting a hierarchical functional organization (101). Whereas feedforward activity is associated with middle cortical layers, feedback connectivity links with superficial and deeper layers. Importantly, there may be key differences between layer-wise connectivity in primary and transmodal areas (102, 103). It has been suggested that, in humans, superficial layers of the transmodal cortex have been implicated in manipulating information, whereas deeper layers may have a more controlling function (104, 105). Interestingly, using high-resolution data it has been shown that frontal regions dominate feedback processes, whereas parietal regions support feedforward functions (106). Such differences may relate to the differential pattern in anterior regions associated with emotion processing, showing somewhat distinct patterning relative to posterior regions across mental-training content. Moreover, various studies in non-human mammals have indicated differential mechanisms of plasticity correspond to different layer-depths (107–109), further underscoring the relevance to consider depth-dependent variation in cortical microstructure over time. It is of note that the current study used a proxy of layer-depth that doesn’t correspond to actual cortical layers, but rather creates depth-varying profiles of 1mm resolution quantitative qT1 though interpolation. However, dept-variations in such profiles have been used to quantify individual differences in both histological sections (110) and *in vivo* during development (93). Future work, incorporating multi-level metrics of cortical morphology using ultra-high-resolution imaging with (mental training) longitudinal design may help to further disentangle the different modes of plasticity in structure and function.

In sum, combining a longitudinal mental training study with multi-modal imaging, we could show that mental training modules focusing on attention, socio-emotional and socio-cognitive skills resulted in differentiable change in intrinsic functional and microstructural organization. In line with prior work revealing differential changes in grey matter morphology after each of the three *ReSource* training modules in the same sample (55), the current work differentiates processes related to our ability of understanding the thoughts and feelings of ourselves and others within the intrinsic functional and microstructural organization of the human brain, providing a potential system-level perspective on social functioning. Although our work focused on healthy adults ranging from 20 to 55 years of age, our findings overall support the possibility that targeted mental training can enhance social skills and lead to co-occurring reconfigurations of cortical function and microstructure, providing evidence for experiencedependent plasticity.

## Materials and Methods

### Experimental design

#### Participants

A total of 332 healthy adults (197 women, mean±SD=40.7±9.2 years, 20-55 years), recruited in 2012-2014 participated in the study, see Table 1 for more details. More than 95% of our sample was Caucasian, with catchment areas balanced across two German municipalities (Berlin and Leipzig). Participant eligibility was determined through a multistage procedure that involved several screening and mental health questionnaires, together with a phone interview [for details, see (54)]. Next, a face-to-face mental health diagnostic interview with a trained clinical psychologist was scheduled. The interview included a computer-assisted German version of the Structured Clinical Interview for DSM-IV Axis-I disorders, SCID-I DIA-X (111) and a personal interview, SCID-II, for Axis-II disorders (112, 113). Participants were excluded if they fulfilled criteria for: i) an Axis-I disorder within the past two years; ii) Schizophrenia, psychotic disorders, bipolar disorder, substance dependency, or an Axis-II disorder at any time in their life. No participant had a history of neurological disorders or head trauma, based on an in-house self-report questionnaire used to screen all volunteers prior to imaging investigations. In addition, participants underwent a diagnostic radiological evaluation to rule out the presence of mass lesions (e.g., tumors, vascular malformations). All participants gave written informed consent and the study was approved by the Research Ethics Committees of the University of Leipzig (#376/12-ff) and Humboldt University in Berlin (#2013-02, 2013-29, 2014-10). The study was registered at ClinicalTrials.gov under the title “Plasticity of the Compassionate Brain” (#NCT01833104). For details on recruitment and sample selection, see the full cohort and study descriptor (54).

#### Sample size estimation and group allocation

Overall, 2595 people signed up for the *ReSource* study in winter 2012/2013. Of these individuals, 311 potential participants met all eligibility criteria. From the latter group, 198 were randomly selected as the final sample. Participants were selected from the larger pool of potential participants and assigned to cohorts using bootstrapping without replacement, creating cohorts that did not differ (omnibus test p<0.1) in demographics (age, gender, marital status, income, and IQ) or self-reported traits (depression, empathy, interoceptive awareness, stress level, compassion for self and others, alexithymia, general mental health, anxiety, agreeableness, conscientiousness, extraversion, neuroticism, and openness). Seven participants dropped out of the study after assignment but before data collection began, leaving 30 participants in RCC1, 80 in TC1, and 81 in TC2.

2144 people applied for the second wave of the study in winter 2013/2014. Of these people, 248 potential participants met all the eligibility criteria. From the latter pool, 164 were then randomly selected as the final sample. Participants were selected from the larger pool of potential participants and assigned to cohorts using bootstrapping without replacement, creating cohorts that did not differ significantly (omnibus test, p>0.1) from the Winter 2012/2013 cohorts or from one another in demographics (age, gender, marital status, income, and IQ) or self-reported traits (depression, empathy, interoceptive awareness, stress level, compassion for self and others, alexithymia, general mental health, anxiety, agreeableness, conscientiousness, neuroticism, and openness). The control cohorts (RCC1, RCC2, and RCC1&2) were significantly lower in extraversion than TC3; participants in the control cohorts were also more likely to have children than participants in TC3. Twenty-three participants dropped out of the study after assignment but before data collection began, leaving 81 participants in TC3 and 60 in RCC2. See further (54).

#### Study design

Our study focused on two training groups: training cohort 1 (TC1, n=80 at enrolment) and training cohort 2 (TC2, n=81 at enrolment), as well as a retest control cohort (RCC) that was partly measured prior to (n=30 at enrolment) and partly after (n=60 at enrolment) TC1 and TC2. A third training cohort (TC3, n=81 at enrolment) underwent an independent training program, and was included as an additional active control for the *Presence* module. Participants were selected from a larger pool of potential volunteers by bootstrapping without replacement, creating cohorts not differing significantly with respect to several demographic and self-report traits (54). Total training duration of TC1 and TC2 was 39 weeks (~nine months), divided into three modules: *Presence, Affect*, and *Perspective* (see below, for details), each lasting for 13 weeks (**Figure 1**). TC3 only participated in one 13-week *Affect* training, and is only included in supplementary robustness analyses, so that the main analysis of functional plasticity focusses on TC1 and TC2. Our main cohorts of interest, TC1 and TC2, underwent *Affect* and *Perspective* modules in different order to act as active control cohorts for each other. Specifically, TC1 underwent “*Presence-Affect-Perspective*”, whereas TC2 underwent “*Presence-Perspective-Affect*”. TC1, TC2, and RCC underwent four testing phases. The baseline-testing phase is called T0; testing phases at the end of the xth module are called Tx (*i.e*., T1, T2, T3). In RCC, testing was carried out at similarly spaced intervals. The study had a slightly time-shifted design, where different groups started at different time points to simultaneously accommodate scanner and teacher availability. As we focused on training-related effects, we did not include analysis of a follow-up measurement T4 that was carried out 4, 5, or 10 months after the official training had ended. For details on training and practice set-up, timeline, and measures, see (54).

#### Final sample

We excluded individuals with missing structural and/or functional MRI data and/or a framewise-displacement of >0.3mm (<5%) (114). Further details of sample size per time-point and exclusion criteria are in **Table 1–3**.

### Neuroimaging acquisition and analysis

#### MRI acquisition

MRI data were acquired on a 3T Siemens Magnetom Verio (Siemens Healthcare, Erlangen, Germany) using a 32-channel head coil. We recorded task-free functional MRI using a T2*-weighted gradient 2D-EPI sequence (repetition time [TR] =2000ms, echo time [TE]=27ms, flip angle=90°; 37 slices tilted at approximately 30° with 3 mm slice thickness, field of view [FOV]=210×210mm2, matrix=70×70, 3×3×3 mm3 voxels, 1 mm gap; 210 volumes per session). We also acquired a T1-weighted 3D-MPRAGE sequence (176 sagittal slices, TR=2300 ms, TE=2.98 ms, inversion time [TI]=900 ms, flip angle=7°, FOV=240×256 mm^2^, matrix=240×256, 1×1×1 mm^3^ voxels). Throughout the duration of our longitudinal study, imaging hardware and console software (Syngo B17) were held constant. For qT1 mapping, we used the recently introduced 3D MP2RAGE sequence (56), which combines two MPRAGE datasets acquired at different inversion times to provide intrinsic bias field cancellation and estimation of T1 (TR=5000 ms, TE=2.89 ms, TI=700/2500 ms, flip angle=4/5°, FOV=256×256 mm2, 1 × 1 × 1–mm3 voxels.

#### Task-free functional MRI analysis

Processing was based on DPARSF/REST for Matlab [http://www.restfmri.net (115)]. We discarded the first 5 volumes to ensure steady-state magnetization, performed slice-time correction, motion correction and realignment, and co-registered functional time series of a given subject to the corresponding T1-weighted MRI. Images underwent unified segmentation and registration to MNI152, followed by nuisance covariate regression to remove effects of average WM and CSF signal, as well as 6 motion parameters (3 translations, 3 rotations). We included a scrubbing (114) that modelled time points with a frame-wise displacement of ≥0.5 mm, together with the preceding and subsequent time points as separate regressors during nuisance covariate correction. Volume-based timeseries were mapped to the fsaverage5 surface using bbregister.

#### Gradient construction

In line with previous studies evaluating functional gradients (32, 36, 62, 73, 83, 116) the functional connectivity matrix was proportionally thresholded at 90% per row and converted into a normalised angle matrix using the BrainSpace toolbox for MATLAB (62). Diffusion map embedding, a nonlinear manifold learning technique (63), identified principal gradient components, explaining functional connectivity variance in descending order. In brief, the algorithm estimates a low-dimensional embedding from a highdimensional affinity matrix. In this space, cortical nodes that are strongly interconnected by either many suprathreshold edges or few very strong edges are closer together, whereas nodes with little or no functional connectivity are farther apart. The name of this approach, which belongs to the family of graph Laplacians, derives from the equivalence of the Euclidean distance between points in the embedded space and the diffusion distance between probability distributions centred at those points. It is controlled by the parameter α, which controls the influence of the density of sampling points on the manifold (α = 0, maximal influence; α = 1, no influence). Based on previous work (36), we set α = 0.5, a choice that retains the global relations between data points in the embedded space and has been suggested to be relatively robust to noise in the functional connectivity matrix. The diffusion time (t), which controls the scale of eigenvalues of the diffusion operator was set at t=0 (default). Individual embedding solutions were aligned to the group-level embedding based on the Human Connectome Project S1200 sample (66) via Procrustes rotations (62). The Procrustes alignment enables comparison across individual embedding solutions, provided the original data is equivalent enough to produce comparable Euclidean spaces.

#### 3D gradient metric: eccentricity

To construct the combined gradient, we computed the Euclidean distance to the individual center for gradient 1-3. Next, to evaluate change within and between individuals, we computed the difference between gradient scores between different time-points.

### Processing of microstructural data

T1-weighted MRIs were processed using FreeSurfer (http://surfer.nmr.mgh.harvard.edu) version 5.1.0 to generate cortical surface models for measurements of cortical thickness and surface area. FreeSurfer has been validated against histological analysis (117) and manual measurements (118). We chose the most general cross-sectional image processing procedure to enable the longitudinal as well as cross sectional study goals of the ReSource Project (see for example (119)). Since data acquisition spanned more than 2 years, a non-specific imaging procedure enabled baseline data analysis before the completion of latter time points, without compromising comparability. Quantitative T1 maps were aligned and mapped to the T1-weighted MRI using a boundary-based registration (120), by maximizing the intensity gradient across tissues boundaries and using the surfaces that separate brain structure and tissue types of the T1-weighted reference image, and the tissue intensity of the quantitative T1 map. Following we computed a dept-dependent microstructure proxy. Depth-dependent cortical microstructure analysis has a long tradition in neuroanatomy (121, 122), and depth-dependent shift in cellular and myelin characteristics have been shown to reflect architectural complexity (122) and cortical hierarchy (38). We generated 12 equivolumetric surfaces between outer and inner cortical surfaces. The equivolumetric model adjusts for cortical folding by varying the Euclidean distance ρ between pairs of intracortical surfaces throughout the cortex to preserve the fractional volume between surfaces. ρ was calculated for each surface (1):

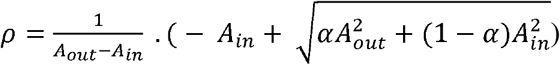

in which α represents a fraction of the total volume of the segment accounted for by the surface, while A_out_ and A_in_ represents the surface area of outer and inner cortical surfaces, respectively. We systematically sampled T1q values for each of the 20,484 linked vertices from the outer to the inner surface across the cortex. Following, we computed the average value of T1q in each of the 400 parcels (67).

### Meta-analytical networks

We used the NeuroSynth meta-analytic database (http://www.neurosynth.org) (61) to assess topic terms associated with the training (“attention”, “interoception”, “emotion”, “empathy”, “Theory of Mind”).

### Behavioral markers

We assessed a battery of behavioral markers developed and adapted to target the main aims of the *Presence, Perspective*, and *Affect* modules: selective attention, compassion, and ToM. Behavioral changes of these markers elicited by the different modules are reported elsewhere (72). The measure for compassion was based on the EmpaToM task, a developed and validated naturalistic video paradigm in the current subjects (91, 123). Videos showed people recounting autobiographical episodes that were either emotionally negative (e.g., loss of a loved one) or neutral (e.g., commuting to work), followed by Likert-scale ratings of experienced valence and compassion. Since the conceptual understanding of compassion might change due to the training, we ensured a consistent understanding by defining it prior to each measurement as experiencing feelings of care, warmth, and benevolence. Compassion was quantified as mean of compassion ratings across all experimental conditions. The EmpaToM task (91) also allowed for measurement of ToM performance. After the ratings, multiple-choice questions requiring inference of mental states (thoughts, intentions, beliefs) of the person in the video or factual reasoning on the video’s content (control condition) were asked. Questions had three response options and only one correct answer, which had been validated during pre-study piloting (91). Here, we calculated participants’ error rates during the ToM questions after the video, collapsed across neutral and negative conditions.

### Statistical analysis

Analysis was performed using SurfStat for Matlab (124). We employed linear mixed-effects models, a flexible statistical technique that allows for inclusion of multiple measurements per subjects and irregular measurement intervals (125). In all models, we controlled for age and sex, and random effect of subject. Inference was performed on subject-specific eccentricity/gradient change maps, Δeccentricity/gradient, which were generated by subtracting vertex-wise eccentricity/gradient maps of subsequent time points for a given participant.

a. *Assessing module-specific change*. To compare the modules against each other, we contrasted change of one given module against the average of the other two modules. To compare two modules, we compared one training module against another (for example *Affect* versus *Perspective*). To compare a given module against RCC, we estimated contrasts for training cohort change relative to RCC (*Presence, Perspective, Affect*).
b. *Correction for multiple comparisons*. Findings were corrected for number of tests within the analysis step using FDR correction (126).

Theoretically, the cross-over design of the study and the inclusion of number of scans since baseline as covariance controlled for test-retest effects on motion (as participants may become calmer in scanner after repeated sessions). Nevertheless, to control for outliers, we removed all individuals with >0.3 mm/degree movement (114).

### Behavioral prediction

We adopted a supervised framework with cross-validation to predict behavioral change based on change in functional and microstructural organization within five functional networks. We aimed at predicting attention, compassion, and perspective-taking (**Fig. 4**). Before running our model, we regressed out age and sex from the brain markers. We utilized 5-fold crossvalidation separating training and test data and repeated this procedure 100 time with different set of training and test data to avoid bias for separating subjects. Following we performed an elastic net cross validation procedure with alphas varying from 0.0001-1, ratio 1.0, making it a lasso regression. We used sequential feature selection to determine the top 20% of features based on mean absolute error without cross validation. Linear regression for predicting behavioral scores was constructed using the selected features as independent variables within the training data (4/5 segments) and it was applied to the test data (1/5 segment) to predict their behavioral scores. The prediction accuracy was assessed by calculating Pearson’s correlation between the actual and predicted behavioral scores as well as their negative mean absolute error, nMAE.

## Acknowledgements

Data for the *ReSource* project were collected between 2013 and 2016 at the Department of Social Neuroscience at the Max Planck Institute for Human Cognitive and Brain Sciences Leipzig. Tania Singer (Principal Investigator) received funding for the ReSource Project from the European Research Council (ERC) under the European Community’s Seventh Framework Program (FP7/2007–2013) ERC grant agreement number 205557. Sofie Valk received support from the Max Planck Society (Otto Hahn Award). Boris Bernhardt acknowledges research support from the NSERC (Discovery-1304413), the Canadian Institutes of Health Research (CIHR FDN-154298), SickKids Foundation (NI17-039), Azrieli Center for Autism Research (ACAR-TACC), and the Tier-2 Canada Research Chairs program. Bo-yong Park was funded by Molson Neuro-Engineering fellowship by Montreal Neurological Institute and Hospital (MNI) and the Fonds de la Recherche du Quebec – Santé. We thank Lisa Feldman Barrett for fruitful discussions during the conception of this manuscript.

## Author contributions

SLV and BCB were involved in data acquisition and processing, and conceived and designed the resting-state computational experiments. SH and BP contributed to development of gradient and genetic enrichment analysis. F.-M.T., A.B., and P.K. designed and analyzed the functional and behavioral data used in this study. T.S. initiated and developed the *ReSource* Project and model, as well as the training protocol. All authors discussed, wrote, and approved the final version of the manuscript.

## Data and code availability

In line with EU data regulations (General Data Protection Regulation, GDPR), we regret that data cannot be shared publicly because we did not obtain explicit participant agreement for data-sharing with third parties. Our work is based on personal data (age, sex and neuroimaging data) that could be matched to individuals. The data is therefore pseudonominized rather than anonymized and falls under the GDPR. Data are available upon request (contact via). Summary data and analysis scripts (Matlab and python) to reproduce primary analyses and figures are publicly available on GitHub (https://github.com/CNG-LAB/social_function_structure_change), and raw data-plots are provided for network-level analyses.

