## Supplementary Materials for "Functional and microstructural plasticity following social and interoceptive mental training"

* joint co-authors

*1. Otto Hahn Group Cognitive Neurogenetics, Max Planck Institute for Human Cognitive and Brain Sciences, Leipzig, Germany; 2. INM-7, FZ Jülich, Jülich, Germany; 3. Clinical Psychology and Behavioral Neuroscience, Faculty of Psychology, Technische Universität Dresden, Dresden, Germany; 4. Max Planck Institute for Human Cognitive and Brain Sciences, Leipzig, Germany; 5. Multimodal Imaging and Connectome Analysis Lab, McConnell Brain Imaging Centre, Montreal Neurological Institute and Hospital, McGill University, Montreal, Quebec, Canada; 6. Department of Data Science, Inha University, Incheon, South Korea; 7. Center for Neuroscience Imaging Research, Institute for Basic Science, Suwon, South Korea; 8. Center for the Developing Brain, Child Mind Institute, NY, USA; 9. Department of Biomedical Engineering, Sungkyunkwan University, Suwon, South Korea; 10. Department of Psychology, Wurzburg University, Germany; 11. Department of Psychosomatic Medicine and Psychotherapy, Medical Center – University of Freiburg, Faculty of Medicine, University of Freiburg, Freiburg im Breisgau, Germany; 12. Social Neuroscience Lab, Max Planck Society, Berlin, Germany*

### Supplementary Tables

**Supplementary Table 1. Descriptive statistics Presence**

| **Presence** | Mean | Std | CI low | CI high |
|---|---|---|---|---|
| **Attention** | 0,002 | 0,013 | -0,001 | 0,004 |
| **Interoception** | 0,003 | 0,015 | 0,000 | 0,006 |
| **Emotion** | 0,001 | 0,012 | -0,002 | 0,003 |
| **Empathy** | 0,002 | 0,011 | 0,000 | 0,004 |
| **ToM** | 0,003 | 0,014 | 0,000 | 0,006 |

**Supplementary Table 2a. Descriptive statistics Affect**

| **Affect** | Mean | Std | CI low | CI high |
|---|---|---|---|---|
| **Attention** | -0,001 | 0,012 | -0,003 | 0,001 |
| **Interoception** | 0,001 | 0,015 | -0,001 | 0,003 |
| **Emotion** | 0,001 | 0,013 | -0,001 | 0,003 |
| **Empathy** | 0,001 | 0,011 | -0,001 | 0,003 |
| **ToM** | 0,000 | 0,016 | -0,002 | 0,003 |

**Supplementary Table 2b. Descriptive statistics Affect, excluding active controls (TC3)**

| **Affect** | Mean | Std | CI low | CI high |
|---|---|---|---|---|
| **Attention** | -0,001 | 0,013 | -0,003 | 0,002 |
| **Interoception** | 0,000 | 0,015 | -0,003 | 0,003 |
| **Emotion** | 0,000 | 0,013 | -0,002 | 0,003 |
| **Empathy** | 0,000 | 0,012 | -0,002 | 0,002 |
| **ToM** | -0,001 | 0,016 | -0,004 | 0,002 |

**Supplementary Table 3. Descriptive statistics Perspective**

| **Perspective** | Mean | Std | CI low | CI high |
|---|---|---|---|---|
| **Attention** | -0,003 | 0,013 | -0,006 | -0,001 |
| **Interoception** | -0,003 | 0,016 | -0,006 | 0,000 |
| **Emotion** | 0,000 | 0,015 | -0,003 | 0,003 |
| **Empathy** | -0,001 | 0,012 | -0,004 | 0,001 |
| **ToM** | -0,001 | 0,018 | -0,005 | 0,003 |

**Supplementary Table 4. Descriptive statistics Retest controls**

| **Retest controls** | Mean | Std | CI low | CI high |
|---|---|---|---|---|
| **Attention** | -0,001 | 0,014 | -0,003 | 0,001 |
| **Interoception** | 0,001 | 0,014 | -0,001 | 0,003 |
| **Emotion** | 0,001 | 0,013 | -0,001 | 0,002 |
| **Empathy** | 0,001 | 0,011 | -0,001 | 0,002 |
| **ToM** | 0,001 | 0,016 | -0,002 | 0,003 |

**Supplementary Table 5. Functional eccentricity changes GSR controlled per functional network.** T-values and p-values below p<0.05, * indicates FDRp<0.05.

| | **Presence vs Perspective** | | **Presence vs Affect** | | **Perspective vs Affect** | |
|---|---|---|---|---|---|---|
| **Attention** | 3,541 | 0,001* | 2,243 | 0,025 | -1,777 | |
| **Interoception** | 3,203 | 0,002* | 1,552 | | -2060 | 0,040 |
| **Emotion** | 1,371 | | 0,892 | | -0,665 | |
| **Empathy** | 2,898 | 0,004* | 1,660 | | -1,621 | |
| **ToM** | 2,047 | 0,041 | 1,449 | | -0,883 | |

**Supplementary Table 6. Functional eccentricity changes in training cohort 1 per functional network.** T-values and p-values below p<0.05, * indicates FDRp<0.05.

| | Presence vs Perspective | | Presence vs Affect | | Perspective vs Affect |
|---|---|---|---|---|---|
| **Attention** | 1,953 | | 1,707 | | -0,245 |
| **Interoception** | 2,332 | 0,021 | 0,889 | | -1,392 |
| **Emotion** | 1,347 | | 0,819 | | -0,512 |
| **Empathy** | 1,984 | 0,049 | 0,950 | | -0,100 |
| **ToM** | 1,908 | | 1,825 | | -0,089 |

**Supplementary Table 7. Functional eccentricity changes in training cohort 2 per functional network.** T-values and p-values below p<0.05, * indicates FDRp<0.05.

| | Presence vs Perspective | | Presence vs Affect | | Perspective vs Affect |
|---|---|---|---|---|---|
| **Attention** | 2,028 | 0,044 | 0,010 | | -1,966 |
| **Interoception** | 1,383 | | 0,794 | | -0,627 |
| **Emotion** | -1,016 | | -0,623 | | 0,421 |
| **Empathy** | 0,906 | | 0,514 | | -0,416 |
| **ToM** | 0,386 | | 0,547 | | 0,144 |

**Supplementary Table 8. Functional eccentricity changes per functional network baseline to T1.** T-values and p-values below p<0.05, * indicates FDRp<0.05.

| | TC1-RCC | | TC2-RCC | | TC3-RCC | | TC1-TC3 | | TC2-TC3 |
|---|---|---|---|---|---|---|---|---|---|
| **Attention** | 2,529 | 0,012 | 1,523 | | 1,144 | | 1,424 | | 0,440 |
| **Interoception** | -0,260 | | -0,813 | | -0,756 | | 0,505 | | -0,093 |
| **Emotion** | 0,411 | | -1,059 | | 0,492 | | -0,077 | | -1,553 |
| **Empathy** | 0,608 | | 0,039 | | 0,348 | | 0,269 | | -0,296 |
| **ToM** | 1,225 | | 0,358 | | 0,763 | | 0,476 | | -0,379 |

**Supplementary Table 9. Functional eccentricity changes per functional network T1 to T3.** T-values and p-values below p<0.05, * indicates FDRp<0.05.

| | Perspective vs Retest Control | | Affect vs Retest Control | | Perspective-Affect |
|---|---|---|---|---|---|
| **Attention** | -1,981 | 0,048 | -0,431 | | -1,528 |
| **Interoception** | -0,960 | | 0,544 | | -1,461 |
| **Emotion** | -0,143 | | -0,045 | | -0,097 |
| **Empathy** | -1,192 | | -0,113 | | -1,059 |
| **ToM** | -0,818 | | -0,858 | | 0,022 |

**Supplementary Table 10. G1-G3 change per functional network Presence vs Perspective.** T-values and p-values below p<0.05, * indicates FDRp<0.05.

| | G1 | | G2 | | G3 |
|---|---|---|---|---|---|
| **Attention** | -1,321 | | 2,508 | 0.012 | 1,098 |
| **Interoception** | -1,787 | | -2,180 | 0.030 | -1,499 |
| **Emotion** | -0,118 | | 1,501 | | 0,833 |
| **Empathy** | -0,573 | | 1,713 | | -0,206 |
| **ToM** | 1,048 | | 2,102 | 0.036 | -0,947 |

**Supplementary Table 11. G1-G3 change per functional network Presence vs Affect.** T-values and p-values below p<0.05, * indicates FDRp<0.05.

| | G1 | | G2 | | G3 |
|---|---|---|---|---|---|
| **Attention** | -0.945 | | 1.715 | | 0,113 |
| **Interoception** | -2.185 | 0.03 | -0.901 | | -1,339 |
| **Emotion** | -0.643 | | 0.871 | | -0,782 |
| **Empathy** | -0.976 | | 3.215 | 0.002* | -1,824 |
| **ToM** | 1.135 | | 2.382 | 0.018 | -0,925 |

**Supplementary Table 12. G1-G3 change per functional network Perspective vs Affect.** T-values and p-values below p<0.05, * indicates FDRp<0.05.

| | G1 | | G2 | | G3 |
|---|---|---|---|---|---|
| **Attention** | 0.527 | | -1.081 | | -1,087 |
| **Interoception** | -0.156 | | 1.506 | | 0,344 |
| **Emotion** | -0.490 | | -0.796 | | -1,654 |
| **Empathy** | -0.315 | | 1.228 | | -1,531 |
| **ToM** | -0.050 | | 0.002 | | 0,140 |

**Supplementary Table 13. Functional eccentricity changes per functional network controlling for cortical thickness change.** T-values and p-values below p<0.05, * indicates FDRp<0.05.

| | Presence vs Perspective | | Presence vs Affect | | Perspective vs Affect | |
|---|---|---|---|---|---|---|
| **Attention** | 2,859 | 0,005* | 1,479 | | -1,691 | |
| **Interoception** | 2,744 | 0,007* | 1,064 | | -1,964 | 0.050 |
| **Emotion** | 0,374 | | -0,165 | | -0,566 | |
| **Empathy** | 2,206 | 0,028 | 0,870 | | -1,566 | |
| **ToM** | 1,709 | | 1,316 | | -0,595 | |

**Supplementary Table 14. Functional eccentricity changes per functional network from baseline to T3.** T-values and p-values below p<0.05, * indicates FDRp<0.05.

| | Training vs Retest Control |
|---|---|
| **Attention** | -0,567 |
| **Interoception** | -0,380 |
| **Emotion** | -0,299 |
| **Empathy** | -0,651 |
| **ToM** | -0,192 |

**Supplementary Table 15. Descriptive of retest-control change (mean change over T0-T1; T1-T2; T2-T3)**

| Control | Attention | | | | Interoception | | | | Emotion | | | |
|---|---|---|---|---|---|---|---|---|---|---|---|---|
| | Mean | Std | CI min | CI max | Mean | Std | CI min | CI max | Mean | Std | CI min | CI max |
| **1** | 7,562 | 49,980 | -0,050 | 15,175 | 1,997 | 32,385 | -2,936 | 6,929 | -2,210 | 28,341 | -6,527 | 2,107 |
| **2** | 7,081 | 40,984 | 0,838 | 13,323 | 1,862 | 25,168 | -1,971 | 5,696 | -2,224 | 21,211 | -5,455 | 1,007 |
| **3** | 6,366 | 34,091 | 1,173 | 11,559 | 1,737 | 20,467 | -1,381 | 4,854 | -2,101 | 16,432 | -4,604 | 0,402 |
| **4** | 5,510 | 28,803 | 1,123 | 9,897 | 1,535 | 17,499 | -1,130 | 4,201 | -1,910 | 13,395 | -3,950 | 0,130 |
| **5** | 4,594 | 24,717 | 0,829 | 8,359 | 1,267 | 15,458 | -1,087 | 3,622 | -1,647 | 11,497 | -3,399 | 0,104 |

| | | | | | | | | | | | | |
|---|---|---|---|---|---|---|---|---|---|---|---|---|
| **6** | 3,718 | 21,721 | 0,409 | 7,026 | 0,933 | 13,867 | -1,180 | 3,045 | -1,350 | 10,542 | -2,955 | 0,256 |
| **7** | 2,951 | 19,821 | -0,069 | 5,970 | 0,546 | 12,560 | -1,367 | 2,460 | -1,069 | 10,405 | -2,654 | 0,516 |
| **8** | 2,357 | 19,003 | -0,537 | 5,252 | 0,159 | 11,639 | -1,614 | 1,932 | -0,844 | 10,843 | -2,495 | 0,808 |
| **9** | 1,987 | 19,123 | -0,925 | 4,900 | -0,199 | 11,126 | -1,894 | 1,496 | -0,697 | 11,507 | -2,450 | 1,056 |
| **10** | 1,841 | 19,858 | -1,184 | 4,865 | -0,529 | 10,911 | -2,191 | 1,133 | -0,630 | 12,042 | -2,464 | 1,204 |
| **11** | 1,891 | 20,844 | -1,284 | 5,066 | -0,831 | 10,900 | -2,491 | 0,829 | -0,670 | 12,163 | -2,522 | 1,183 |
| **12** | 2,088 | 21,799 | -1,232 | 5,409 | -1,073 | 10,979 | -2,745 | 0,600 | -0,814 | 11,806 | -2,612 | 0,984 |

| | **Empathy** | | | | **ToM** | | | |
|---|---|---|---|---|---|---|---|---|
| **1** | 2,651 | 30,526 | -1,999 | 7,301 | 1,606 | 25,823 | -2,328 | 5,539 |
| **2** | 1,865 | 23,063 | -1,648 | 5,378 | 1,080 | 19,190 | -1,844 | 4,003 |
| **3** | 1,270 | 17,727 | -1,430 | 3,970 | 0,669 | 14,902 | -1,601 | 2,938 |
| **4** | 0,828 | 14,162 | -1,329 | 2,985 | 0,383 | 12,308 | -1,492 | 2,258 |
| **5** | 0,531 | 11,871 | -1,278 | 2,339 | 0,207 | 10,757 | -1,431 | 1,846 |
| **6** | 0,343 | 10,550 | -1,264 | 1,950 | 0,121 | 9,905 | -1,388 | 1,629 |
| **7** | 0,237 | 10,039 | -1,292 | 1,766 | 0,090 | 9,681 | -1,385 | 1,565 |
| **8** | 0,171 | 10,209 | -1,384 | 1,726 | 0,077 | 10,020 | -1,449 | 1,604 |
| **9** | 0,123 | 10,749 | -1,514 | 1,761 | 0,057 | 10,617 | -1,560 | 1,674 |
| **10** | 0,061 | 11,323 | -1,664 | 1,786 | -0,001 | 11,112 | -1,694 | 1,692 |
| **11** | -0,034 | 11,723 | -1,819 | 1,752 | -0,112 | 11,273 | -1,829 | 1,605 |
| **12** | -0,160 | 11,866 | -1,967 | 1,647 | -0,274 | 11,079 | -1,962 | 1,413 |

**Supplementary Table 16. Descriptive of Presence change (mean change over T0-T1)**

| | **Attention** | | | | **Interoception** | | | | **Emotion** | |
|---|---|---|---|---|---|---|---|---|---|---|
| | Mean | Std | CI min | CI max | Mean | Std | CI min | CI max | Mean | Std |
| **1** | 26,962 | 42,445 | 18,904 | 35,021 | 11,744 | 32,692 | 5,5366 | 17,95 | -0,1614 | 29,164 |
| **2** | 22,432 | 37,77 | 15,261 | 29,602 | 10,628 | 24,888 | 5,9029 | 15,35 | -0,75176 | 21,216 |
| **3** | 17,804 | 34,339 | 11,284 | 24,324 | 9,0319 | 20,491 | 5,1414 | 12,92 | -1,2386 | 17,084 |
| **4** | 13,436 | 31,073 | 7,5364 | 19,335 | 7,2546 | 18,145 | 3,8096 | 10,7 | -1,5975 | 14,796 |
| **5** | 9,636 | 27,622 | 4,3918 | 14,88 | 5,4632 | 16,527 | 2,3254 | 8,601 | -1,8056 | 13,27 |
| **6** | 6,6193 | 24,229 | 2,0193 | 11,219 | 3,7751 | 15,091 | 0,90993 | 6,64 | -1,8841 | 12,369 |
| **7** | 4,4032 | 21,348 | 0,35014 | 8,4563 | 2,3068 | 13,884 | -0,32914 | 4,943 | -1,8585 | 12,251 |
| **8** | 2,9186 | 19,435 | -0,77129 | 6,6085 | 1,0881 | 13,073 | -1,3938 | 3,57 | -1,8242 | 12,83 |
| **9** | 2,048 | 18,648 | -1,4924 | 5,5884 | 0,09958 | 12,75 | -2,3212 | 2,52 | -1,8667 | 13,723 |
| **10** | 1,6593 | 18,743 | -1,8993 | 5,2179 | -0,69095 | 12,736 | -3,1091 | 1,727 | -2,0169 | 14,379 |
| **11** | 1,5937 | 19,274 | -2,0655 | 5,253 | -1,3077 | 12,618 | -3,7032 | 1,088 | -2,2674 | 14,323 |
| **12** | 1,732 | 19,852 | -2,0371 | 5,5011 | -1,7709 | 12,094 | -4,0669 | 0,525 | -2,5839 | 13,256 |

| | CI min | CI max | **Empathy** | | | | **ToM** | | | |
|---|---|---|---|---|---|---|---|---|---|---|
| **1** | -5,6984 | 5,3756 | 10,435 | 28,632 | 4,9986 | 15,87 | 8,5313 | 22,468 | 4,2655 | 12,8 |
| **2** | -4,7798 | 3,2763 | 8,126 | 22,669 | 3,8221 | 12,43 | 5,8231 | 16,579 | 2,6755 | 8,971 |
| **3** | -4,4821 | 2,0049 | 6,1404 | 19,553 | 2,4281 | 9,853 | 3,7308 | 14,063 | 1,0608 | 6,401 |
| **4** | -4,4067 | 1,2117 | 4,506 | 17,422 | 1,1984 | 7,814 | 2,1832 | 13,019 | -0,28858 | 4,655 |
| **5** | -4,3251 | 0,71385 | 3,2537 | 15,371 | 0,33528 | 6,172 | 1,1235 | 12,101 | -1,1739 | 3,421 |
| **6** | -4,2325 | 0,46425 | 2,39 | 13,492 | -0,17144 | 4,952 | 0,53595 | 11,18 | -1,5866 | 2,659 |

| | | | | | | | | | | |
|---|---|---|---|---|---|---|---|---|---|---|
| **7** | -4,1844 | 0,46747 | 1,7933 | 12,311 | -0,54403 | 4,131 | 0,30078 | 10,795 | -1,7487 | 2,35 |
| **8** | -4,2601 | 0,61178 | 1,3569 | 12,164 | -0,95258 | 3,666 | 0,26041 | 11,319 | -1,8886 | 2,409 |
| **9** | -4,4722 | 0,73882 | 0,9688 | 12,865 | -1,4737 | 3,411 | 0,27816 | 12,455 | -2,0866 | 2,643 |
| **10** | -4,7468 | 0,71295 | 0,591 | 13,757 | -2,0208 | 3,203 | 0,25582 | 13,511 | -2,3094 | 2,821 |
| **11** | -4,9868 | 0,45196 | 0,2205 | 14,239 | -2,4828 | 2,924 | 0,12787 | 13,9 | -2,5111 | 2,767 |
| **12** | -5,1007 | -0,06723 | -0,136 | 14,066 | -2,8062 | 2,535 | -0,1157 | 13,375 | -2,655 | 2,424 |

**Supplementary Table 17a. Descriptive of Affect change (mean change over T0-T1, T1-T2 and T2-T3)**

| | **Attention** | | | | **Interoception** | | | | **Emotion** | |
|---|---|---|---|---|---|---|---|---|---|---|
| | Mean | Std | CI min | CI max | Mean | Std | CI min | CI max | Mean | Std |
| **1** | 4,706 | 44,870 | -2,212 | 11,625 | 1,224 | 29,272 | -3,289 | 5,738 | -6,308 | 25,848 |
| **2** | 2,206 | 38,832 | -3,782 | 8,193 | 0,772 | 23,371 | -2,832 | 4,376 | -5,746 | 19,466 |
| **3** | 0,288 | 33,868 | -4,934 | 5,510 | 0,107 | 19,716 | -2,933 | 3,147 | -5,139 | 15,474 |
| **4** | -1,147 | 29,495 | -5,695 | 3,401 | -0,647 | 17,393 | -3,329 | 2,035 | -4,647 | 12,938 |
| **5** | -2,203 | 25,537 | -6,140 | 1,735 | -1,357 | 15,627 | -3,767 | 1,053 | -4,306 | 11,271 |
| **6** | -2,943 | 22,076 | -6,347 | 0,461 | -1,969 | 14,056 | -4,137 | 0,198 | -4,102 | 10,373 |
| **7** | -3,446 | 19,395 | -6,437 | -0,456 | -2,498 | 12,669 | -4,451 | -0,544 | -4,024 | 10,246 |
| **8** | -3,791 | 17,745 | -6,528 | -1,055 | -2,982 | 11,649 | -4,778 | -1,186 | -4,065 | 10,788 |
| **9** | -4,057 | 17,136 | -6,700 | -1,415 | -3,421 | 11,112 | -5,134 | -1,707 | -4,187 | 11,688 |
| **10** | -4,303 | 17,292 | -6,969 | -1,637 | -3,799 | 10,982 | -5,493 | -2,106 | -4,350 | 12,497 |
| **11** | -4,529 | 17,818 | -7,276 | -1,781 | -4,111 | 11,001 | -5,807 | -2,415 | -4,471 | 12,750 |
| **12** | -4,699 | 18,383 | -7,534 | -1,865 | -4,300 | 10,904 | -5,982 | -2,619 | -4,496 | 12,191 |
| | CI min | CI max | **Empathy** | | | | **ToM** | | | |
| **1** | -10,293 | -2,322 | -0,781 | 26,852 | -4,921 | 3,359 | -0,289 | 20,940 | -3,517 | 2,940 |
| **2** | -8,748 | -2,745 | -1,624 | 21,451 | -4,931 | 1,684 | -1,155 | 17,108 | -3,793 | 1,483 |
| **3** | -7,525 | -2,753 | -2,234 | 17,936 | -5,000 | 0,531 | -1,769 | 14,871 | -4,062 | 0,524 |
| **4** | -6,642 | -2,652 | -2,629 | 15,423 | -5,007 | -0,251 | -2,190 | 13,319 | -4,243 | -0,136 |
| **5** | -6,044 | -2,568 | -2,889 | 13,394 | -4,954 | -0,824 | -2,449 | 12,020 | -4,303 | -0,596 |
| **6** | -5,702 | -2,503 | -3,073 | 11,807 | -4,894 | -1,253 | -2,576 | 11,004 | -4,273 | -0,880 |
| **7** | -5,604 | -2,445 | -3,224 | 10,873 | -4,901 | -1,548 | -2,618 | 10,536 | -4,242 | -0,993 |
| **8** | -5,729 | -2,402 | -3,366 | 10,721 | -5,019 | -1,713 | -2,621 | 10,736 | -4,276 | -0,965 |
| **9** | -5,989 | -2,385 | -3,529 | 11,195 | -5,255 | -1,803 | -2,625 | 11,397 | -4,383 | -0,868 |
| **10** | -6,277 | -2,423 | -3,714 | 11,910 | -5,550 | -1,877 | -2,659 | 12,160 | -4,534 | -0,784 |
| **11** | -6,437 | -2,505 | -3,911 | 12,458 | -5,832 | -1,990 | -2,731 | 12,680 | -4,686 | -0,776 |
| **12** | -6,376 | -2,617 | -4,114 | 12,614 | -6,059 | -2,169 | -2,843 | 12,745 | -4,809 | -0,878 |

**Supplementary Table 17b. Descriptive of Affect change (mean change over T1-T2 and T2-T3)**

| | **Attention** | | | | **Interoception** | | | | **Emotion** | |
|---|---|---|---|---|---|---|---|---|---|---|
| | Mean | Std | CI min | CI max | Mean | Std | CI min | CI max | Mean | Std |
| **1** | -2,5228 | 46,28 | -11,523 | 6,4775 | -1,6892 | 30,662 | -7,6521 | 4,274 | -5,2022 | 26,077 |
| **2** | -4,0713 | 39,93 | -11,837 | 3,6942 | -2,287 | 24,126 | -6,9789 | 2,405 | -4,9854 | 19,189 |
| **3** | -4,9563 | 34,624 | -11,69 | 1,7773 | -2,8899 | 19,931 | -6,766 | 0,986 | -4,6893 | 14,886 |
| **4** | -5,3472 | 29,986 | -11,179 | 0,4843 | -3,416 | 17,237 | -6,7682 | -0,064 | -4,4435 | 12,235 |

| | | | | | | | | | | |
|---|---|---|---|---|---|---|---|---|---|---|
| **5** | -5,4408 | 25,904 | -10,478 | -0,4032 | -3,7642 | 15,357 | -6,7508 | -0,778 | -4,2724 | 10,623 |
| **6** | -5,3062 | 22,483 | -9,6786 | -0,9339 | -3,9564 | 13,899 | -6,6593 | -1,254 | -4,1187 | 9,9775 |
| **7** | -5,063 | 19,975 | -8,9476 | -1,1785 | -4,0454 | 12,751 | -6,5251 | -1,566 | -4,0143 | 10,208 |
| **8** | -4,7844 | 18,592 | -8,4001 | -1,1687 | -4,1359 | 11,948 | -6,4596 | -1,812 | -3,9849 | 11,065 |
| **9** | -4,5494 | 18,277 | -8,1038 | -0,995 | -4,2382 | 11,532 | -6,4808 | -1,996 | -3,9986 | 12,15 |
| **10** | -4,4092 | 18,688 | -8,0435 | -0,7749 | -4,3449 | 11,422 | -6,5663 | -2,124 | -4,0351 | 12,983 |
| **11** | -4,3492 | 19,414 | -8,1248 | -0,5736 | -4,4493 | 11,388 | -6,664 | -2,235 | -4,0192 | 13,092 |
| **12** | -4,3433 | 20,116 | -8,2554 | -0,4312 | -4,49 | 11,182 | -6,6646 | -2,316 | -3,8985 | 12,234 |

| | CI min | CI max | **Empathy** | | | | **ToM** | | | |
|---|---|---|---|---|---|---|---|---|---|---|
| **1** | -10,273 | -0,13092 | -3,236 | 27,882 | -8,6579 | 2,187 | -2,019 | 20,485 | -6,0027 | 1,965 |
| **2** | -8,7171 | -1,2537 | -4,01 | 22,161 | -8,3195 | 0,3 | -2,7088 | 16,61 | -5,9391 | 0,522 |
| **3** | -7,5841 | -1,7944 | -4,491 | 18,239 | -8,0381 | -0,944 | -3,1489 | 14,324 | -5,9345 | -0,363 |
| **4** | -6,8228 | -2,0641 | -4,685 | 15,376 | -7,6756 | -1,695 | -3,4087 | 12,741 | -5,8864 | -0,931 |
| **5** | -6,3384 | -2,2065 | -4,688 | 13,145 | -7,2448 | -2,132 | -3,5069 | 11,45 | -5,7338 | -1,28 |
| **6** | -6,0591 | -2,1783 | -4,599 | 11,56 | -6,8469 | -2,351 | -3,4807 | 10,514 | -5,5254 | -1,436 |
| **7** | -5,9995 | -2,029 | -4,473 | 10,79 | -6,5717 | -2,375 | -3,3644 | 10,184 | -5,3449 | -1,384 |
| **8** | -6,1367 | -1,8331 | -4,35 | 10,863 | -6,4623 | -2,237 | -3,2093 | 10,526 | -5,2563 | -1,162 |
| **9** | -6,3615 | -1,6357 | -4,263 | 11,539 | -6,5068 | -2,019 | -3,0571 | 11,297 | -5,254 | -0,86 |
| **10** | -6,56 | -1,5102 | -4,233 | 12,413 | -6,6467 | -1,819 | -2,932 | 12,161 | -5,297 | -0,567 |
| **11** | -6,5652 | -1,4731 | -4,244 | 13,099 | -6,7913 | -1,696 | -2,8149 | 12,79 | -5,3022 | -0,328 |
| **12** | -6,2778 | -1,5192 | -4,286 | 13,409 | -6,8935 | -1,678 | -2,7091 | 12,996 | -5,2365 | -0,182 |

**Supplementary Table 18. Descriptive of Perspective change (mean change over T1-T2 and T2-T3)**

| | **Attention** | | | | **Interoception** | | | | **Emotion** | |
|---|---|---|---|---|---|---|---|---|---|---|
| | Mean | Std | CI min | CI max | Mean | Std | CI min | CI max | Mean | Std |
| **1** | -10,87 | 52,435 | -21,494 | -0,2458 | -2,7462 | 34,112 | -9,6579 | 4,165 | -2,7201 | 27,766 |
| **2** | -9,2606 | 44,418 | -18,26 | -0,2608 | -3,2496 | 27,916 | -8,9059 | 2,407 | -2,9624 | 21,214 |
| **3** | -7,5768 | 38,091 | -15,295 | 0,1412 | -3,371 | 23,751 | -8,1835 | 1,442 | -2,6939 | 17,138 |
| **4** | -5,9606 | 32,896 | -12,626 | 0,7047 | -3,2017 | 20,677 | -7,3913 | 0,988 | -2,237 | 14,413 |
| **5** | -4,5253 | 28,585 | -10,317 | 1,2667 | -2,8753 | 18,163 | -6,5554 | 0,805 | -1,7131 | 12,627 |
| **6** | -3,3834 | 25,12 | -8,4732 | 1,7064 | -2,4586 | 16,021 | -5,7048 | 0,788 | -1,2131 | 11,73 |
| **7** | -2,534 | 22,66 | -7,1253 | 2,0573 | -1,9479 | 14,33 | -4,8514 | 0,956 | -0,74685 | 11,697 |
| **8** | -1,947 | 21,333 | -6,2694 | 2,3755 | -1,3666 | 13,157 | -4,0326 | 1,299 | -0,35126 | 12,308 |
| **9** | -1,58 | 21,038 | -5,8427 | 2,6827 | -0,79743 | 12,511 | -3,3324 | 1,738 | -0,045665 | 13,176 |
| **10** | -1,3974 | 21,448 | -5,7432 | 2,9483 | -0,28119 | 12,209 | -2,7549 | 2,193 | 0,17334 | 13,835 |
| **11** | -1,3603 | 22,168 | -5,8519 | 3,1313 | 0,14272 | 12,018 | -2,2924 | 2,578 | 0,21532 | 13,883 |
| **12** | -1,4199 | 22,878 | -6,0554 | 3,2156 | 0,45214 | 11,769 | -1,9325 | 2,837 | 0,083819 | 13,132 |

| | CI min | CI max | **Empathy** | | | | **ToM** | | | |
|---|---|---|---|---|---|---|---|---|---|---|
| **1** | -8,3461 | 2,9059 | -4,22 | 32,346 | -10,774 | 2,334 | -4,5243 | 23,152 | -9,2154 | 0,167 |
| **2** | -7,2608 | 1,336 | -3,914 | 25,651 | -9,1112 | 1,284 | -3,8974 | 18,304 | -7,6062 | -0,189 |
| **3** | -6,1663 | 0,77853 | -3,437 | 20,779 | -7,6477 | 0,773 | -3,1972 | 15,084 | -6,2534 | -0,141 |
| **4** | -5,1574 | 0,68333 | -2,87 | 17,117 | -6,3379 | 0,599 | -2,5082 | 12,903 | -5,1227 | 0,106 |

| | | | | | | | | | | |
|---|---|---|---|---|---|---|---|---|---|---|
| **5** | -4,2715 | 0,84524 | -2,307 | 14,48 | -5,2405 | 0,627 | -1,9118 | 11,426 | -4,227 | 0,403 |
| **6** | -3,5899 | 1,1637 | -1,828 | 12,794 | -4,4202 | 0,765 | -1,443 | 10,636 | -3,598 | 0,712 |
| **7** | -3,1169 | 1,6232 | -1,422 | 12,087 | -3,8706 | 1,027 | -1,1308 | 10,646 | -3,2879 | 1,026 |
| **8** | -2,8451 | 2,1426 | -1,113 | 12,185 | -3,5823 | 1,356 | -0,94722 | 11,328 | -3,2424 | 1,348 |
| **9** | -2,7153 | 2,624 | -0,893 | 12,736 | -3,474 | 1,687 | -0,83875 | 12,296 | -3,3301 | 1,653 |
| **10** | -2,6299 | 2,9766 | -0,758 | 13,363 | -3,4655 | 1,95 | -0,79372 | 13,216 | -3,4715 | 1,884 |
| **11** | -2,5977 | 3,0283 | -0,722 | 13,757 | -3,5091 | 2,066 | -0,81567 | 13,77 | -3,6058 | 1,975 |
| **12** | -2,577 | 2,7447 | -0,759 | 13,798 | -3,5551 | 2,036 | -0,89601 | 13,851 | -3,7024 | 1,91 |

**Supplementary Table 19. Layer T1q change per functional network Presence vs Perspective.** T-values and p-values below p<0.05, and Cohen's D effect size, * indicates FDRp<0.05.

| | **Attention** | | | **Interoception** | | | **Emotion** | | | **Empathy** | | | **ToM** | | |
|---|---|---|---|---|---|---|---|---|---|---|---|---|---|---|---|
| | t | p | D | t- | p | D | t | p | D | t | p | D | t | p | D |
| **1** | 5,663 | 0,001* | 0,491 | 3,217 | 0,001* | 0,279 | 0,639 | | 0,055 | 3,539 | 0,001* | 0,307 | 3,992 | 0,001* | 0,346 |
| **2** | 5,581 | 0,001* | 0,484 | 3,914 | 0,001* | 0,340 | 0,753 | | 0,065 | 3,725 | 0,001* | 0,323 | 3,868 | 0,001* | 0,336 |
| **3** | 5,177 | 0,001* | 0,449 | 4,207 | 0,001* | 0,365 | 0,630 | | 0,055 | 3,642 | 0,001* | 0,316 | 3,349 | 0,001* | 0,291 |
| **4** | 4,552 | 0,001* | 0,395 | 4,065 | 0,001* | 0,353 | 0,332 | | 0,029 | 3,330 | 0,001* | 0,289 | 2,604 | 0,009* | 0,226 |
| **5** | 3,816 | 0,001* | 0,331 | 3,629 | 0,001* | 0,315 | -0,056 | | -0,005 | 2,920 | 0,004* | 0,253 | 1,878 | | 0,163 |
| **6** | 3,075 | 0,002* | 0,267 | 3,014 | 0,003* | 0,262 | -0,434 | | -0,038 | 2,508 | 0,012* | 0,218 | 1,324 | | 0,115 |
| **7** | 2,376 | 0,018* | 0,206 | 2,262 | 0,024 | 0,196 | -0,729 | | -0,063 | 2,042 | 0,042 | 0,177 | 0,977 | | 0,085 |
| **8** | 1,778 | | 0,154 | 1,394 | | 0,121 | -0,934 | | -0,081 | 1,560 | | 0,135 | 0,786 | | 0,068 |
| **9** | 1,344 | | 0,117 | 0,511 | | 0,044 | -1,085 | | -0,094 | 1,109 | | 0,096 | 0,673 | | 0,058 |
| **10** | 1,109 | | 0,096 | -0,280 | | -0,024 | -1,240 | | -0,108 | 0,749 | | 0,065 | 0,591 | | 0,051 |
| **11** | 1,037 | | 0,090 | -0,916 | | -0,080 | -1,394 | | -0,121 | 0,495 | | 0,043 | 0,515 | | 0,045 |
| **12** | 1,075 | | 0,093 | -1,414 | | -0,123 | -1,570 | | -0,136 | 0,321 | | 0,028 | 0,434 | | 0,038 |

**Supplementary Table 20. Layer T1q change per functional network Presence vs Affect.** T-values and p-values below p<0.05, and Cohen's D effect size, * indicates FDRp<0.05.

| | **Attention** | | | **Interoception** | | | **Emotion** | | | **Empathy** | | | **ToM** | | |
|---|---|---|---|---|---|---|---|---|---|---|---|---|---|---|---|
| | t | p | D | t- | p | D | t | p | D | t | p | D | t | p | D |
| **1** | 3,840 | 0,001* | 0,333 | 2,722 | 0,007* | 0,236 | 1,800 | | 0,156 | 3,082 | 0,002* | 0,267 | 3,081 | 0,002* | 0,267 |
| **2** | 4,094 | 0,001* | 0,355 | 3,243 | 0,001* | 0,281 | 1,940 | | 0,168 | 3,419 | 0,001* | 0,297 | 3,167 | 0,002* | 0,275 |
| **3** | 4,101 | 0,001* | 0,356 | 3,536 | 0,000* | 0,307 | 1,914 | | 0,166 | 3,592 | 0,001* | 0,312 | 3,019 | 0,003* | 0,262 |
| **4** | 3,928 | 0,001* | 0,341 | 3,591 | 0,000* | 0,312 | 1,793 | | 0,156 | 3,640 | 0,001* | 0,316 | 2,750 | 0,006* | 0,239 |
| **5** | 3,671 | 0,001* | 0,319 | 3,478 | 0,001* | 0,302 | 1,680 | | 0,146 | 3,647 | 0,001* | 0,317 | 2,504 | 0,013* | 0,217 |
| **6** | 3,396 | 0,001* | 0,295 | 3,266 | 0,001* | 0,283 | 1,605 | | 0,139 | 3,671 | 0,001* | 0,319 | 2,372 | 0,018* | 0,206 |
| **7** | 3,122 | 0,002* | 0,271 | 3,015 | 0,003* | 0,262 | 1,575 | | 0,137 | 3,636 | 0,001* | 0,316 | 2,293 | 0,022 | 0,199 |
| **8** | 2,860 | 0,004* | 0,248 | 2,743 | 0,006* | 0,238 | 1,579 | | 0,137 | 3,436 | 0,001* | 0,298 | 2,182 | 0,030 | 0,189 |

| | | | | | | | | | | | | | | | |
|---|---|---|---|---|---|---|---|---|---|---|---|---|---|---|---|
| **9** | 2,641 | 0,009* | 0,229 | 2,450 | 0,015* | 0,213 | 1,540 | | 0,134 | 3,119 | 0,002* | 0,271 | 2,045 | 0,041 | 0,177 |
| **10** | 2,512 | 0,012* | 0,218 | 2,163 | 0,031 | 0,188 | 1,478 | | 0,128 | 2,816 | 0,005* | 0,244 | 1,919 | | 0,167 |
| **11** | 2,468 | 0,014* | 0,214 | 1,932 | | 0,168 | 1,393 | | 0,121 | 2,602 | 0,010* | 0,226 | 1,818 | | 0,158 |
| **12** | 2,482 | 0,013* | 0,215 | 1,774 | | 0,154 | 1,277 | | 0,111 | 2,488 | 0,013* | 0,216 | 1,742 | | 0,151 |

**Supplementary Table 21. Layer T1q change per functional network Perspective vs Affect.** T-values and p-values below p<0.05, and Cohen's D effect size, * indicates FDRp<0.05.

| | **Attention** | | | **Interoception** | | | **Emotion** | | | **Empathy** | | | **ToM** | | |
|---|---|---|---|---|---|---|---|---|---|---|---|---|---|---|---|
| | t | p | D | t- | p | D | t | p | D | t | p | D | t | p | D |
| **1** | -2,455 | 0,014 | -0,213 | -0,876 | | -0,076 | 1,035 | | 0,090 | -0,883 | | -0,077 | -1,377 | | -0,120 |
| **2** | -2,122 | 0,034 | -0,184 | -1,131 | | -0,098 | 1,046 | | 0,091 | -0,763 | | -0,066 | -1,160 | | -0,101 |
| **3** | -1,679 | | -0,146 | -1,167 | | -0,101 | 1,155 | | 0,100 | -0,511 | | -0,044 | -0,742 | | -0,064 |
| **4** | -1,166 | | -0,101 | -0,959 | | -0,083 | 1,364 | | 0,118 | -0,125 | | -0,011 | -0,191 | | -0,017 |
| **5** | -0,615 | | -0,053 | -0,595 | | -0,052 | 1,676 | | 0,146 | 0,328 | | 0,028 | 0,363 | | 0,032 |
| **6** | -0,075 | | -0,006 | -0,131 | | -0,011 | 2,017 | 0,044 | 0,175 | 0,800 | | 0,069 | 0,841 | | 0,073 |
| **7** | 0,421 | | 0,037 | 0,443 | | 0,038 | 2,309 | 0,021 | 0,200 | 1,275 | | 0,111 | 1,142 | | 0,099 |
| **8** | 0,818 | | 0,071 | 1,122 | | 0,097 | 2,533 | 0,012* | 0,220 | 1,608 | | 0,140 | 1,242 | | 0,108 |
| **9** | 1,079 | | 0,094 | 1,799 | | 0,156 | 2,658 | 0,008* | 0,231 | 1,792 | | 0,156 | 1,234 | | 0,107 |
| **10** | 1,210 | | 0,105 | 2,383 | 0,018* | 0,207 | 2,766 | 0,006* | 0,240 | 1,892 | | 0,164 | 1,202 | | 0,104 |
| **11** | 1,245 | | 0,108 | 2,857 | 0,004* | 0,248 | 2,850 | 0,005* | 0,247 | 1,962 | 0,050 | 0,170 | 1,188 | | 0,103 |
| **12** | 1,217 | | 0,106 | 3,246 | 0,001* | 0,282 | 2,930 | 0,004* | 0,254 | 2,043 | 0,042 | 0,177 | 1,203 | | 0,104 |

**Supplementary Table 22. Layer T1q change per functional network Presence vs Perspective in TC1.** T-values and p-values below p<0.05. * indicates FDRp<0.05.

| | **Attention** | | **Interoception** | | **Emotion** | | **Empathy** | | **ToM** | |
|---|---|---|---|---|---|---|---|---|---|---|
| **1** | 4,136 | 0.001* | 2,618 | 0,010 | 0,568 | | 3,285 | 0,001* | 3,445 | 0,001* |
| **2** | 3,741 | 0,001* | 2,873 | 0,005* | 0,421 | | 3,228 | 0,002* | 2,951 | 0,004* |
| **3** | 3,165 | 0,002* | 2,857 | 0,005* | 0,144 | | 2,922 | 0,004* | 2,152 | 0,033 |
| **4** | 2,553 | 0,012 | 2,599 | 0,010 | -0,166 | | 2,544 | 0,012 | 1,390 | |
| **5** | 1,984 | 0,049 | 2,244 | 0,026 | -0,413 | | 2,254 | 0,026 | 0,897 | |
| **6** | 1,522 | | 1,900 | | -0,535 | | 2,102 | 0,037 | 0,709 | |
| **7** | 1,188 | | 1,556 | | -0,561 | | 1,996 | 0,048 | 0,757 | |
| **8** | 0,983 | | 1,225 | | -0,545 | | 1,871 | | 0,908 | |
| **9** | 0,891 | | 0,883 | | -0,554 | | 1,696 | | 1,019 | |
| **10** | 0,881 | | 0,547 | | -0,601 | | 1,543 | | 1,063 | |
| **11** | 0,910 | | 0,236 | | -0,688 | | 1,425 | | 1,048 | |
| **12** | 0,941 | | -0,061 | | -0,868 | | 1,314 | | 0,964 | |

**Supplementary Table 23. Layer T1q change per functional network Presence vs Affect in TC1.** T-values and p-values below p<0.05. * indicates FDRp<0.05.

| | **Attention** | | **Interoception** | | **Emotion** | | **Empathy** | | **ToM** | |
|---|---|---|---|---|---|---|---|---|---|---|
| **1** | 3,483 | 0,001* | 2,057 | 0,041 | 0,757 | | 2,975 | 0,003* | 2,837 | 0,005* |
| **2** | 3,765 | 0,000* | 2,584 | 0,011 | 0,626 | | 3,330 | 0,001* | 2,687 | 0,008* |

| | | | | | | | | | | |
|---|---|---|---|---|---|---|---|---|---|---|
| **3** | 3,756 | 0,000* | 3,005 | 0,003* | 0,464 | | 3,464 | 0,001* | 2,296 | 0,023 |
| **4** | 3,544 | 0,001* | 3,250 | 0,001* | 0,291 | | 3,439 | 0,001* | 1,821 | 0,071 |
| **5** | 3,226 | 0,002* | 3,326 | 0,001* | 0,156 | | 3,369 | 0,001* | 1,434 | |
| **6** | 2,860 | 0,005* | 3,262 | 0,001* | 0,041 | | 3,278 | 0,001* | 1,189 | |
| **7** | 2,484 | 0,014* | 3,080 | 0,002* | -0,034 | | 3,048 | 0,003* | 1,008 | |
| **8** | 2,126 | 0,035 | 2,811 | 0,006* | -0,083 | | 2,637 | 0,009 | 0,843 | |
| **9** | 1,836 | 0,068 | 2,466 | 0,015* | -0,136 | | 2,159 | 0,032 | 0,696 | |
| **10** | 1,671 | 0,097 | 2,145 | 0,034 | -0,161 | | 1,797 | 0,074 | 0,585 | |
| **11** | 1,619 | | 1,898 | | -0,206 | | 1,571 | | 0,474 | |
| **12** | 1,651 | | 1,737 | | -0,320 | | 1,444 | | 0,354 | |

**Supplementary Table 24. Layer T1q change per functional network Perspective vs Affect in TC1.** T-values and p-values below p<0.05. * indicates FDRp<0.05.

| | **Attention** | | **Interoception** | | **Emotion** | | **Empathy** | | **ToM** |
|---|---|---|---|---|---|---|---|---|---|
| **1** | -0,646 | | -0,550 | | 0,178 | | -0,313 | | -0,599 |
| **2** | 0,004 | | -0,291 | | 0,194 | | 0,081 | | -0,268 |
| **3** | 0,549 | | 0,127 | | 0,305 | | 0,504 | | 0,127 |
| **4** | 0,934 | | 0,610 | | 0,438 | | 0,843 | | 0,405 |
| **5** | 1,178 | | 1,024 | | 0,546 | | 1,055 | | 0,509 |
| **6** | 1,272 | | 1,293 | | 0,554 | | 1,114 | | 0,455 |
| **7** | 1,233 | | 1,449 | | 0,507 | | 0,996 | | 0,236 |
| **8** | 1,088 | | 1,510 | | 0,445 | | 0,723 | | -0,066 |
| **9** | 0,899 | | 1,509 | | 0,403 | | 0,433 | | -0,314 |
| **10** | 0,752 | | 1,525 | | 0,424 | | 0,235 | | -0,463 |
| **11** | 0,673 | | 1,589 | | 0,464 | | 0,132 | | -0,555 |
| **12** | 0,674 | | 1,719 | | 0,529 | | 0,118 | | -0,588 |

**Supplementary Table 25. Layer T1q change per functional network Presence vs Perspective in TC2.** T-values and p-values below p<0.05. * indicates FDRp<0.05.

| | **Attention** | | **Interoception** | | **Emotion** | | **Empathy** | | **ToM** | |
|---|---|---|---|---|---|---|---|---|---|---|
| **1** | 3,891 | 0,001* | 1,765 | 0,080 | 0,304 | | 1,620 | | 2,464 | 0,015 |
| **2** | 4,061 | 0,001* | 2,499 | 0,014 | 0,684 | | 1,895 | | 2,755 | 0,007 |
| **3** | 3,992 | 0,001* | 2,931 | 0,004 | 0,831 | | 1,998 | 0,048 | 2,714 | 0,007 |
| **4** | 3,697 | 0,001* | 3,008 | 0,003 | 0,735 | | 1,903 | | 2,390 | 0,018 |
| **5** | 3,243 | 0,001 | 2,774 | 0,006 | 0,443 | | 1,637 | | 1,890 | |
| **6** | 2,692 | 0,008 | 2,263 | 0,025 | 0,033 | | 1,245 | | 1,308 | |
| **7** | 2,075 | 0,040 | 1,543 | | -0,354 | | 0,727 | | 0,727 | |
| **8** | 1,457 | | 0,658 | | -0,655 | | 0,173 | | 0,233 | |
| **9** | 0,941 | | -0,214 | | -0,866 | | -0,307 | | -0,107 | |
| **10** | 0,610 | | -0,971 | | -1,048 | | -0,679 | | -0,311 | |
| **11** | 0,474 | | -1,532 | | -1,183 | | -0,916 | | -0,415 | |
| **12** | 0,502 | | -1,896 | | -1,251 | | -1,025 | | -0,417 | |

**Supplementary Table 26. Layer T1q change per functional network Presence vs Affect in TC2.** T-values and p-values below p<0.05. * indicates FDRp<0.05.

| | Attention | | Interoception | | Emotion | | Empathy | | ToM | |
|---|---|---|---|---|---|---|---|---|---|---|
| 1 | 2,851 | 0,005 | 2,071 | 0,040 | 1,006 | | 1,617 | | 2,005 | 0,047 |
| 2 | 2,871 | 0,005 | 2,501 | 0,013 | 1,468 | | 1,887 | | 2,424 | 0,017 |
| 3 | 2,783 | 0,006 | 2,668 | 0,008 | 1,746 | | 2,078 | 0,039 | 2,650 | 0,009 |
| 4 | 2,621 | 0,010 | 2,593 | 0,010 | 1,906 | | 2,197 | 0,030 | 2,771 | 0,006 |
| 5 | 2,436 | 0,016 | 2,343 | 0,021 | 2,017 | 0,046 | 2,281 | 0,024 | 2,840 | 0,005 |
| 6 | 2,243 | 0,026 | 2,002 | 0,047 | 2,063 | 0,041 | 2,354 | 0,020 | 2,860 | 0,005 |
| 7 | 2,032 | 0,044 | 1,637 | | 2,061 | 0,041 | 2,353 | 0,020 | 2,758 | 0,007 |
| 8 | 1,793 | | 1,308 | | 2,016 | 0,046 | 2,235 | 0,027 | 2,514 | 0,013 |
| 9 | 1,563 | | 1,046 | | 1,925 | | 2,036 | 0,044 | 2,233 | 0,027 |
| 10 | 1,376 | | 0,813 | | 1,767 | | 1,800 | | 1,976 | |
| 11 | 1,252 | | 0,657 | | 1,579 | | 1,604 | | 1,778 | |
| 12 | 1,192 | | 0,560 | | 1,391 | | 1,484 | | 1,632 | |

**Supplementary Table 27. Layer T1q change per functional network Perspective vs Affect in TC2.** T-values and p-values below $p<0.05$. * indicates $FDRp<0.05$.

| | Attention | | Interoception | | Emotion | | Empathy | | ToM | |
|---|---|---|---|---|---|---|---|---|---|---|
| 1 | -1,156 | | 0,240 | | 0,680 | | -0,059 | | -0,536 | |
| 2 | -1,312 | | -0,085 | | 0,747 | | -0,074 | | -0,420 | |
| 3 | -1,327 | | -0,361 | | 0,870 | | 0,011 | | -0,157 | |
| 4 | -1,187 | | -0,513 | | 1,125 | | 0,223 | | 0,291 | |
| 5 | -0,907 | | -0,521 | | 1,532 | | 0,576 | | 0,868 | |
| 6 | -0,535 | | -0,336 | | 1,993 | 0,048 | 1,047 | | 1,480 | |
| 7 | -0,114 | | 0,039 | | 2,385 | 0,018 | 1,572 | | 1,971 | |
| 8 | 0,280 | | 0,616 | | 2,647 | 0,009 | 2,020 | 0,045 | 2,234 | 0,027 |
| 9 | 0,578 | | 1,245 | | 2,772 | 0,006 | 2,312 | 0,022 | 2,303 | 0,023 |
| 10 | 0,731 | | 1,787 | | 2,801 | 0,006 | 2,459 | 0,015 | 2,258 | 0,025 |
| 11 | 0,748 | | 2,204 | 0,029 | 2,754 | 0,007 | 2,508 | 0,013 | 2,168 | 0,032 |
| 12 | 0,661 | | 2,479 | 0,014 | 2,640 | 0,009 | 2,501 | 0,013 | 2,028 | 0,044 |

**Supplementary Table 28. Layer T1q change per functional network baseline – T1: TC1 (Presence) versus Retest Control.** T-values and p-values below $p<0.05$. * indicates $FDRp<0.05$.

| | Attention | | Interoception | | Emotion | | Empathy | | ToM |
|---|---|---|---|---|---|---|---|---|---|
| 1 | -0,188 | | 0,236 | | 0,691 | | 0,296 | | -0,002 |
| 2 | -0,484 | | 0,082 | | 0,907 | | 0,379 | | -0,316 |
| 3 | -0,764 | | -0,159 | | 0,908 | | 0,286 | | -0,649 |
| 4 | -1,049 | | -0,368 | | 0,691 | | 0,152 | | -0,968 |
| 5 | -1,268 | | -0,507 | | 0,339 | | 0,000 | | -1,259 |
| 6 | -1,401 | | -0,623 | | -0,066 | | -0,132 | | -1,465 |
| 7 | -1,429 | | -0,625 | | -0,409 | | -0,223 | | -1,507 |
| 8 | -1,334 | | -0,530 | | -0,624 | | -0,266 | | -1,348 |
| 9 | -1,152 | | -0,368 | | -0,705 | | -0,306 | | -1,066 |
| 10 | -0,966 | | -0,158 | | -0,684 | | -0,253 | | -0,783 |
| 11 | -0,847 | | 0,053 | | -0,600 | | -0,143 | | -0,525 |

| | | | | | | | | | |
|---|---|---|---|---|---|---|---|---|---|
| **12** | -0,820 | | 0,206 | | -0,454 | | 0,006 | | -0,276 |

**Supplementary Table 29. Layer T1q change per functional network baseline – T1: TC2 (Presence) versus Retest Control.** T-values and p-values below p<0.05. * indicates FDRp<0.05.

| | **Attention** | | **Interoception** | | **Emotion** | | **Empathy** | | **ToM** |
|---|---|---|---|---|---|---|---|---|---|
| **1** | -0,371 | | -0,052 | | 0,467 | | -0,825 | | -0,286 |
| **2** | -0,422 | | 0,209 | | 1,220 | | -0,474 | | 0,119 |
| **3** | -0,487 | | 0,257 | | 1,672 | | -0,259 | | 0,409 |
| **4** | -0,634 | | 0,133 | | 1,766 | | -0,139 | | 0,475 |
| **5** | -0,783 | | -0,094 | | 1,567 | | -0,167 | | 0,327 |
| **6** | -0,935 | | -0,441 | | 1,132 | | -0,324 | | 0,040 |
| **7** | -1,093 | | -0,744 | | 0,605 | | -0,577 | | -0,297 |
| **8** | -1,231 | | -0,985 | | 0,132 | | -0,870 | | -0,578 |
| **9** | -1,319 | | -1,089 | | -0,196 | | -1,145 | | -0,721 |
| **10** | -1,373 | | -1,049 | | -0,393 | | -1,307 | | -0,758 |
| **11** | -1,409 | | -0,885 | | -0,454 | | -1,347 | | -0,686 |
| **12** | -1,439 | | -0,667 | | -0,341 | | -1,264 | | -0,486 |

**Supplementary Table 30. Layer T1q change per functional network baseline – T1: Affect TC3 vs Retest Control.** T-values and p-values below p<0.05. * indicates FDRp<0.05.

| | **Attention** | | **Interoception** | | **Emotion** | | **Empathy** | | **ToM** |
|---|---|---|---|---|---|---|---|---|---|
| **1** | -1,397 | | -0,818 | | -0,896 | | -1,493 | | -1,420 |
| **2** | -1,728 | | -0,881 | | -0,525 | | -1,278 | | -1,382 |
| **3** | -1,948 | | -0,987 | | -0,209 | | -1,031 | | -1,278 |
| **4** | -2,089 | 0,038 | -1,093 | | -0,078 | | -0,903 | | -1,218 |
| **5** | -2,205 | 0,029 | -1,195 | | -0,178 | | -0,968 | | -1,303 |
| **6** | -2,341 | 0,020 | -1,284 | | -0,522 | | -1,227 | | -1,513 |
| **7** | -2,488 | 0,013 | -1,383 | | -0,965 | | -1,614 | | -1,744 |
| **8** | -2,628 | 0,009 | -1,471 | | -1,344 | | -1,957 | | -1,863 |
| **9** | -2,734 | 0,007 | -1,504 | | -1,578 | | -2,134 | 0,034 | -1,835 |
| **10** | -2,821 | 0,005 | -1,446 | | -1,688 | | -2,175 | 0,031 | -1,751 |
| **11** | -2,904 | 0,004 | -1,317 | | -1,701 | | -2,157 | 0,032 | -1,665 |
| **12** | -2,978 | 0,003 | -1,155 | | -1,618 | | -2,118 | 0,035 | -1,578 |

**Supplementary Table 31. Layer T1q change per functional network baseline – T1: Presence TC1 vs Affect TC3.** T-values and p-values below p<0.05. * indicates FDRp<0.05.

| | **Attention** | | **Interoception** | | **Emotion** | | **Empathy** | | **ToM** |
|---|---|---|---|---|---|---|---|---|---|
| **1** | 1,234 | | 1,078 | | 1,624 | | 1,829 | | 1,449 |
| **2** | 1,269 | | 0,984 | | 1,467 | | 1,684 | | 1,088 |
| **3** | 1,207 | | 0,845 | | 1,145 | | 1,326 | | 0,640 |
| **4** | 1,047 | | 0,740 | | 0,788 | | 1,059 | | 0,253 |
| **5** | 0,934 | | 0,698 | | 0,529 | | 0,968 | | 0,041 |
| **6** | 0,932 | | 0,661 | | 0,465 | | 1,097 | | 0,044 |
| **7** | 1,054 | | 0,755 | | 0,567 | | 1,405 | | 0,237 |

| | | | | | | | | | |
|---|---|---|---|---|---|---|---|---|---|
| 8 | 1,297 | | 0,939 | | 0,734 | | 1,726 | | 0,522 |
| 9 | 1,597 | | 1,140 | | 0,890 | | 1,867 | | 0,782 |
| 10 | 1,877 | | 1,301 | | 1,024 | | 1,963 | | 0,987 |
| 11 | 2,077 | 0,039 | 1,389 | | 1,123 | | 2,057 | 0,041 | 1,163 |
| 12 | 2,166 | 0,031 | 1,382 | | 1,187 | | 2,171 | 0,031 | 1,329 |

**Supplementary Table 32. Layer T1q change per functional network baseline – T1: Presence TC2 vs Affect TC3.** T-values and p-values below p<0.05. * indicates FDRp<0.05.

| | Attention | | Interoception | | Emotion | | Empathy | | ToM |
|---|---|---|---|---|---|---|---|---|---|
| 1 | 0,988 | | 0,748 | | 1,356 | | 0,616 | | 1,097 |
| 2 | 1,259 | | 1,077 | | 1,764 | | 0,757 | | 1,475 |
| 3 | 1,408 | | 1,230 | | 1,918 | | 0,725 | | 1,670 |
| 4 | 1,381 | | 1,207 | | 1,886 | | 0,722 | | 1,680 |
| 5 | 1,332 | | 1,070 | | 1,779 | | 0,756 | | 1,611 |
| 6 | 1,306 | | 0,790 | | 1,670 | | 0,848 | | 1,522 |
| 7 | 1,288 | | 0,573 | | 1,565 | | 0,973 | | 1,404 |
| 8 | 1,288 | | 0,412 | | 1,452 | | 1,024 | | 1,231 |
| 9 | 1,308 | | 0,340 | | 1,344 | | 0,916 | | 1,057 |
| 10 | 1,340 | | 0,329 | | 1,250 | | 0,791 | | 0,938 |
| 11 | 1,377 | | 0,374 | | 1,200 | | 0,732 | | 0,927 |
| 12 | 1,405 | | 0,440 | | 1,235 | | 0,778 | | 1,047 |

**Supplementary Table 33. Layer T1q change per functional network T1 – T3: Perspective versus Affect (TC1+TC2).** T-values and p-values below p<0.05. * indicates FDRp<0.05.

| Affect-Control | Attention | | Interoception | | Emotion | | Empathy | | ToM |
|---|---|---|---|---|---|---|---|---|---|
| 1 | -1,175 | | -0,205 | | 0,659 | | -0,202 | | -0,749 |
| 2 | -0,862 | | -0,236 | | 0,720 | | 0,061 | | -0,444 |
| 3 | -0,499 | | -0,133 | | 0,904 | | 0,430 | | 0,004 |
| 4 | -0,112 | | 0,105 | | 1,205 | | 0,867 | | 0,534 |
| 5 | 0,283 | | 0,410 | | 1,598 | | 1,328 | | 1,043 |
| 6 | 0,631 | | 0,754 | | 1,934 | | 1,734 | | 1,434 |
| 7 | 0,901 | | 1,151 | | 2,149 | 0,032 | 2,005 | | 1,592 |
| 8 | 1,057 | | 1,619 | | 2,246 | 0,025 | 2,086 | 0,046 | 1,537 |
| 9 | 1,105 | | 2,080 | 0,038 | 2,270 | 0,024 | 2,051 | 0,038 | 1,401 |
| 10 | 1,084 | | 2,482 | 0,014 | 2,293 | 0,023 | 1,988 | 0,041 | 1,266 |
| 11 | 1,030 | | 2,808 | 0,005 | 2,294 | 0,022 | 1,930 | | 1,143 |
| 12 | 0,968 | | 3,043 | 0,003 | 2,275 | 0,024 | 1,900 | | 1,036 |

**Supplementary Table 34. Layer T1q change per functional network T1 – T3: Affect vs Retest Control. *** T-values and p-values below p<0.05. * indicates FDRp<0.05.

| Affect-Control | Attention | | Interoception | | Emotion | | Empathy | | ToM |
|---|---|---|---|---|---|---|---|---|---|
| 1 | 0,237 | | 0,124 | | -0,569 | | -0,078 | | -0,030 |
| 2 | -0,059 | | 0,105 | | -1,025 | | -0,517 | | -0,495 |
| 3 | -0,363 | | -0,028 | | -1,459 | | -1,010 | | -0,926 |
| 4 | -0,633 | | -0,242 | | -1,798 | | -1,446 | | -1,305 |

| | | | | | | | | | |
|---|---|---|---|---|---|---|---|---|---|
| **5** | -0,858 | | -0,473 | | -2,004 | 0,046 | -1,756 | | -1,544 |
| **6** | -1,007 | | -0,681 | | -2,015 | 0,045 | -1,907 | | -1,624 |
| **7** | -1,072 | | -0,867 | | -1,890 | | -1,871 | | -1,555 |
| **8** | -1,062 | | -1,068 | | -1,735 | | -1,701 | | -1,387 |
| **9** | -1,025 | | -1,279 | | -1,614 | | -1,512 | | -1,238 |
| **10** | -1,000 | | -1,483 | | -1,563 | | -1,389 | | -1,136 |
| **11** | -1,007 | | -1,678 | | -1,563 | | -1,344 | | -1,073 |
| **12** | -1,047 | | -1,858 | | -1,629 | | -1,392 | | -1,084 |

**Supplementary Table 35. Layer T1q change per functional network T1 – T3: Perspective vs Retest Control.** T-values and p-values below p<0.05. * indicates FDRp<0.05.

| | **Attention** | | **Interoception** | | **Emotion** | | **Empathy** | | **ToM** |
|---|---|---|---|---|---|---|---|---|---|
| **1** | -0,937 | | -0,088 | | 0,09 | | -0,276 | | -0,794 |
| **2** | -0,908 | | -0,132 | | -0,293 | | -0,446 | | -0,933 |
| **3** | -0,845 | | -0,159 | | -0,535 | | -0,565 | | -0,903 |
| **4** | -0,729 | | -0,134 | | -0,569 | | -0,561 | | -0,743 |
| **5** | -0,562 | | -0,061 | | -0,382 | | -0,412 | | -0,471 |
| **6** | -0,368 | | 0,0708 | | -0,062 | | -0,162 | | -0,162 |
| **7** | -0,168 | | 0,276 | | 0,273 | | 0,138 | | 0,066 |
| **8** | -0,005 | | 0,537 | | 0,522 | | 0,385 | | 0,183 |
| **9** | 0,079 | | 0,784 | | 0,665 | | 0,536 | | 0,202 |
| **10** | 0,084 | | 0,982 | | 0,738 | | 0,597 | | 0,172 |
| **11** | 0,025 | | 1,117 | | 0,736 | | 0,583 | | 0,110 |
| **12** | -0,074 | | 1,183 | | 0,649 | | 0,505 | | -0,011 |

**Supplementary Table 36. Layer T1q change per functional network controlling for CTX: Presence versus Perspective.** T-values and p-values below p<0.05. * indicates FDRp<0.05.

| | **Attention** | | **Interoception** | | **Emotion** | | **Empathy** | | **ToM** | |
|---|---|---|---|---|---|---|---|---|---|---|
| **1** | 5,664 | 0,001* | 3,264 | 0,001* | 0,425 | | 3,468 | 0,001* | 3,897 | 0,001* |
| **2** | 5,568 | 0,001* | 3,939 | 0,001* | 0,528 | | 3,655 | 0,001* | 3,788 | 0,001* |
| **3** | 5,155 | 0,001* | 4,205 | 0,001* | 0,396 | | 3,568 | 0,001* | 3,278 | 0,001* |
| **4** | 4,520 | 0,001* | 4,049 | 0,001* | 0,082 | | 3,249 | 0,001* | 2,524 | 0,012* |
| **5** | 3,768 | 0,001* | 3,602 | 0,001* | -0,337 | | 2,821 | 0,005* | 1,767 | |
| **6** | 2,996 | 0,003* | 2,962 | 0,003* | -0,759 | | 2,377 | 0,018* | 1,163 | |
| **7** | 2,261 | 0,024 | 2,164 | 0,031 | -1,089 | | 1,881 | | 0,766 | |
| **8** | 1,634 | | 1,240 | | -1,296 | | 1,376 | | 0,544 | |
| **9** | 1,196 | | 0,313 | | -1,437 | | 0,924 | | 0,423 | |
| **10** | 0,984 | | -0,493 | | -1,574 | | 0,577 | | 0,354 | |
| **11** | 0,959 | | -1,128 | | -1,700 | | 0,344 | | 0,308 | |
| **12** | 1,068 | | -1,615 | | -1,833 | | 0,196 | | 0,276 | |

**Supplementary Table 37. Layer T1q change per functional network controlling for CTX: Presence versus Affect.** T-values and p-values below p<0.05. * indicates FDRp<0.05.

| | **Attention** | | **Interoception** | | **Emotion** | | **Empathy** | | **ToM** | |
|---|---|---|---|---|---|---|---|---|---|---|
| **1** | 3,914 | 0,001* | 2,793 | 0,005* | 1,372 | | 3,114 | 0,002* | 2,885 | 0,004* |

| | | | | | | | | | | |
|---|---|---|---|---|---|---|---|---|---|---|
| **2** | 4,140 | 0,001* | 3,274 | 0,001* | 1,534 | | 3,459 | 0,001* | 3,006 | 0,003* |
| **3** | 4,125 | 0,001* | 3,544 | 0,001* | 1,524 | | 3,654 | 0,001* | 2,908 | 0,004* |
| **4** | 3,938 | 0,001* | 3,612 | 0,001* | 1,390 | | 3,716 | 0,001* | 2,680 | 0,008* |
| **5** | 3,669 | 0,001* | 3,540 | 0,001* | 1,230 | | 3,738 | 0,001* | 2,464 | 0,014* |
| **6** | 3,379 | 0,001* | 3,385 | 0,001* | 1,084 | | 3,770 | 0,001* | 2,342 | 0,020 |
| **7** | 3,086 | 0,002* | 3,190 | 0,002* | 0,985 | | 3,704 | 0,001* | 2,243 | 0,025 |
| **8** | 2,806 | 0,005* | 2,959 | 0,003* | 0,916 | | 3,478 | 0,001* | 2,101 | 0,036 |
| **9** | 2,582 | 0,010* | 2,683 | 0,008* | 0,858 | | 3,135 | 0,002* | 1,936 | |
| **10** | 2,463 | 0,014* | 2,388 | 0,017* | 0,806 | | 2,810 | 0,005* | 1,790 | |
| **11** | 2,444 | 0,015* | 2,137 | 0,033 | 0,747 | | 2,574 | 0,010* | 1,678 | |
| **12** | 2,498 | 0,013* | 1,927 | | 0,682 | | 2,439 | 0,015* | 1,602 | |

**Supplementary Table 38. Layer T1q change per functional network controlling for CTX: Perspective versus Affect.** T-values and p-values below $p<0.05$. * indicates FDRp<0.05.

| | **Attention** | | **Interoception** | | **Emotion** | | **Empathy** | | **ToM** |
|---|---|---|---|---|---|---|---|---|---|
| **1** | -2,386 | 0,017* | -0,859 | | 0,857 | | -0,776 | | -1,461 |
| **2** | -2,066 | 0,039 | -1,127 | | 0,901 | | -0,650 | | -1,227 |
| **3** | -1,632 | | -1,157 | | 1,035 | | -0,371 | | -0,770 |
| **4** | -1,122 | | -0,922 | | 1,247 | | 0,037 | | -0,170 |
| **5** | -0,564 | | -0,506 | | 1,549 | | 0,524 | | 0,446 |
| **6** | -0,006 | | 0,040 | | 1,869 | | 1,038 | | 0,987 |
| **7** | 0,511 | | 0,717 | | 2,133 | 0,033 | 1,514 | | 1,324 |
| **8** | 0,923 | | 1,497 | | 2,292 | 0,022 | 1,848 | | 1,428 |
| **9** | 1,182 | | 2,238 | 0,026 | 2,388 | 0,017* | 2,009 | 0,045 | 1,401 |
| **10** | 1,298 | | 2,831 | 0,005* | 2,486 | 0,013* | 2,074 | 0,039 | 1,336 |
| **11** | 1,306 | | 3,283 | 0,001* | 2,564 | 0,011* | 2,100 | 0,036 | 1,278 |
| **12** | 1,240 | | 3,612 | 0,001* | 2,646 | 0,008* | 2,131 | 0,034 | 1,240 |

**Supplementary Table 39. Layer T1q change per functional network Training vs Retest Control.** T-values and p-values below $p<0.05$. * indicates FDRp<0.05.

| | **Attention** | | **Interoception** | | **Emotion** | | **Empathy** | | **ToM** | |
|---|---|---|---|---|---|---|---|---|---|---|
| **1** | -1,368 | | -0,190 | | -0,414 | | -1,099 | | -0,799 | |
| **2** | -2,012 | | -0,454 | | -0,669 | | -1,451 | | -1,338 | |
| **3** | -2,590 | 0,011 | -0,865 | | -1,013 | | -1,807 | | -1,822 | |
| **4** | -3,080 | 0,002* | -1,295 | | -1,441 | | -2,152 | 0,033* | -2,242 | 0,026* |
| **5** | -3,493 | 0,001* | -1,733 | | -1,923 | | -2,531 | 0,012* | -2,617 | 0,010* |
| **6** | -3,820 | 0,001* | -2,164 | 0,032* | -2,367 | 0,019* | -2,958 | 0,004* | -2,976 | 0,003* |
| **7** | -4,035 | 0,001* | -2,512 | 0,013* | -2,615 | 0,010* | -3,340 | 0,001* | -3,249 | 0,001* |
| **8** | -4,090 | 0,001* | -2,754 | 0,007* | -2,622 | 0,010* | -3,523 | 0,001* | -3,262 | 0,001* |
| **9** | -3,987 | 0,001* | -2,859 | 0,005* | -2,481 | 0,014* | -3,497 | 0,001* | -3,023 | 0,003* |
| **10** | -3,819 | 0,001* | -2,769 | 0,006* | -2,360 | 0,020* | -3,388 | 0,001* | -2,733 | 0,007* |
| **11** | -3,694 | 0,001* | -2,551 | 0,012* | -2,324 | 0,021* | -3,303 | 0,001* | -2,482 | 0,014* |
| **12** | -3,659 | 0,001* | -2,288 | 0,023* | -2,363 | 0,019* | -3,256 | 0,001* | -2,282 | 0,024* |

### Supplementary Figures

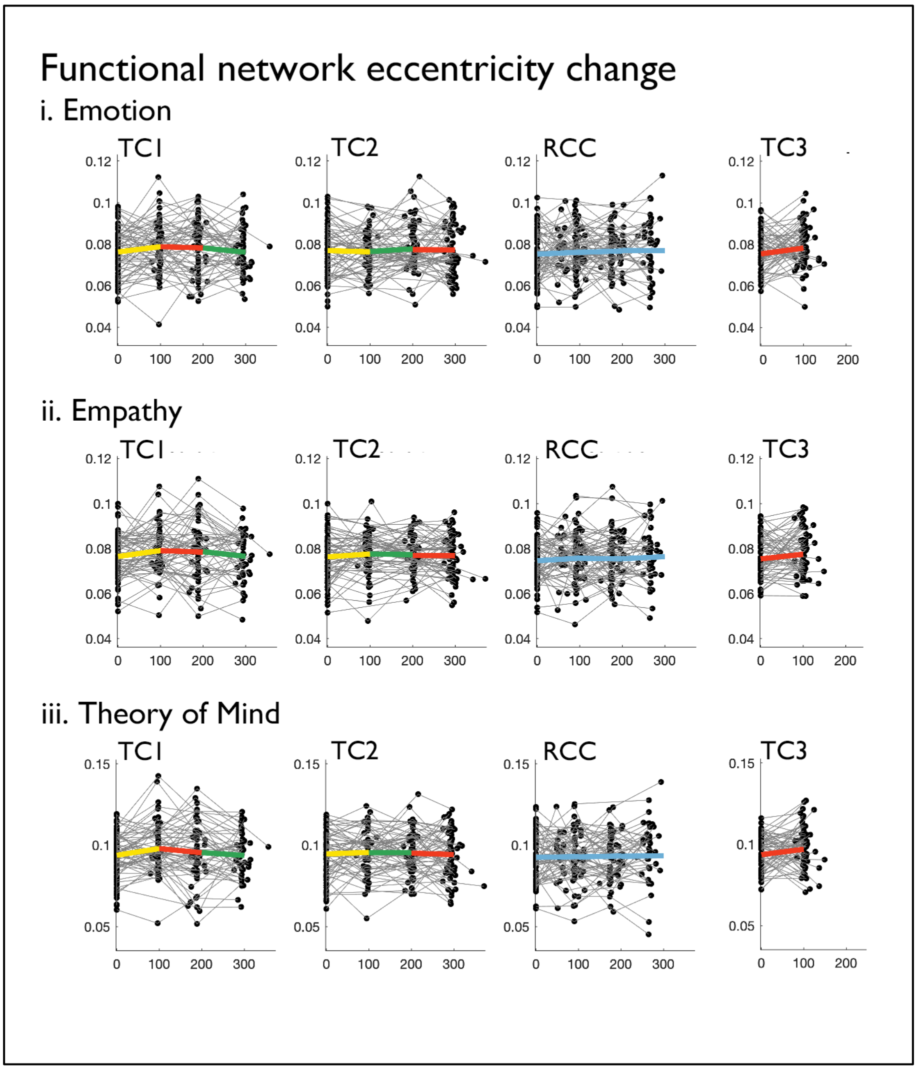

**Supplementary Figure 1. Module-specific change in functional eccentricity as a function of training cohort and functional network.**

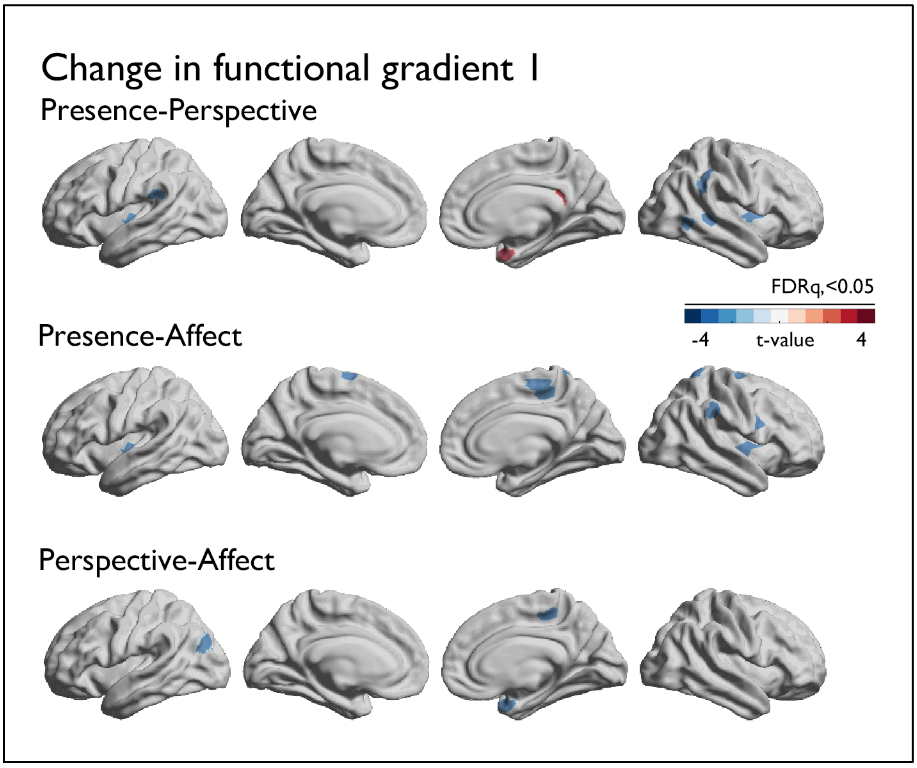

**Supplementary Figure 2. Module-specific change in functional Gradient 1.** Trends at p<0.01, FDRq<0.05 outlined in black.

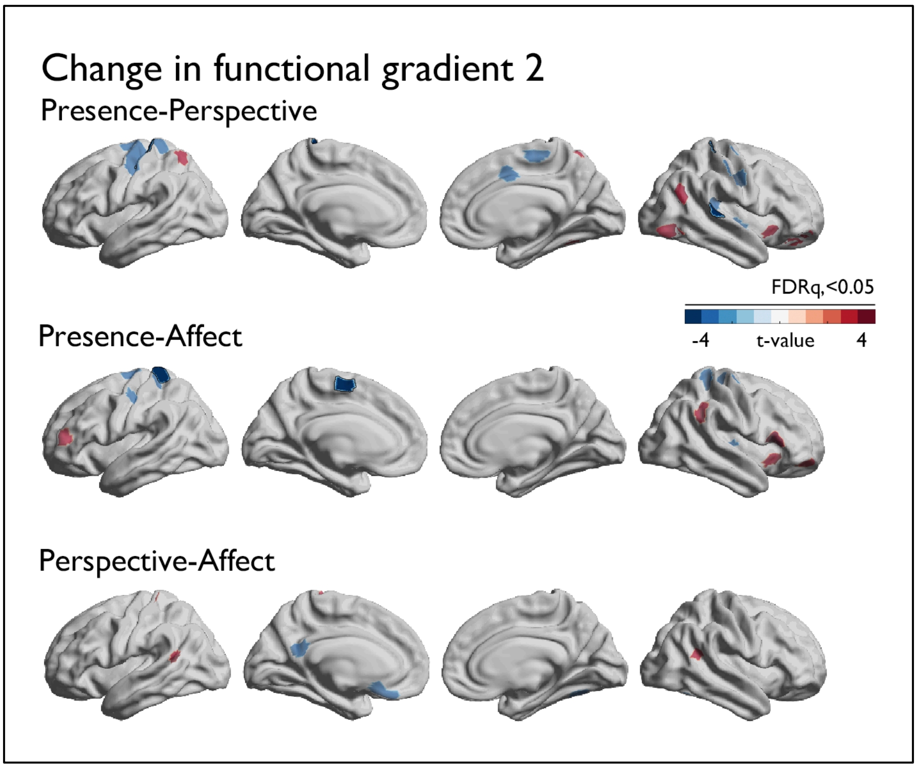

**Supplementary Figure 3. Module-specific change in functional Gradient 2.** Trends at p<0.01, FDRq<0.05 outlined in black.

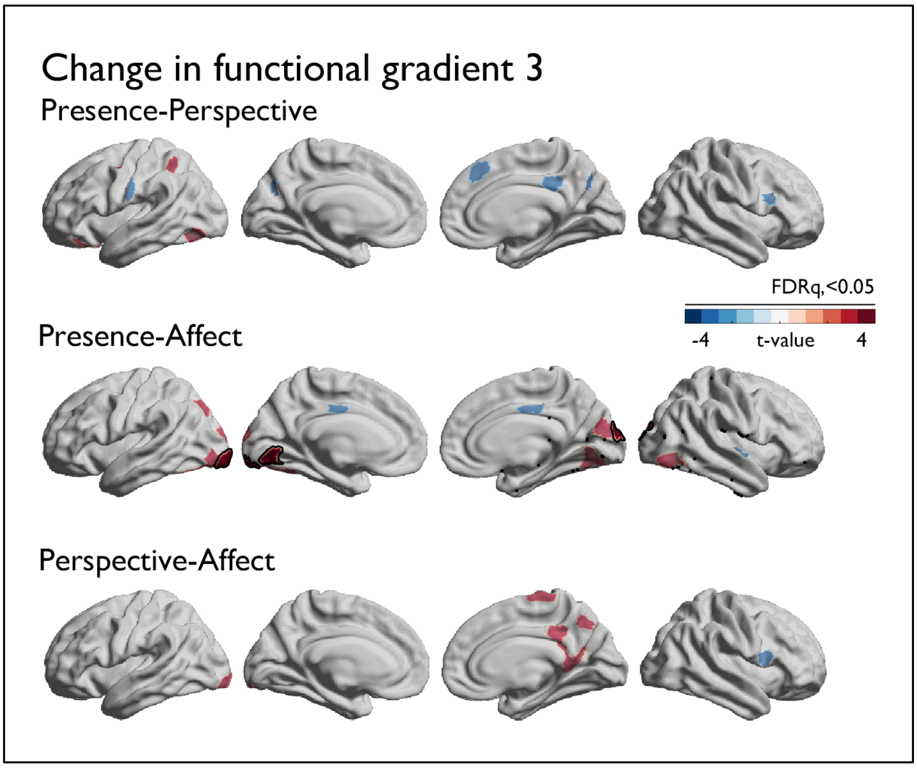

**Supplementary Figure 4. Module-specific change in functional Gradient 3.** Trends at p<0.01, FDRq<0.05 outlined in black.

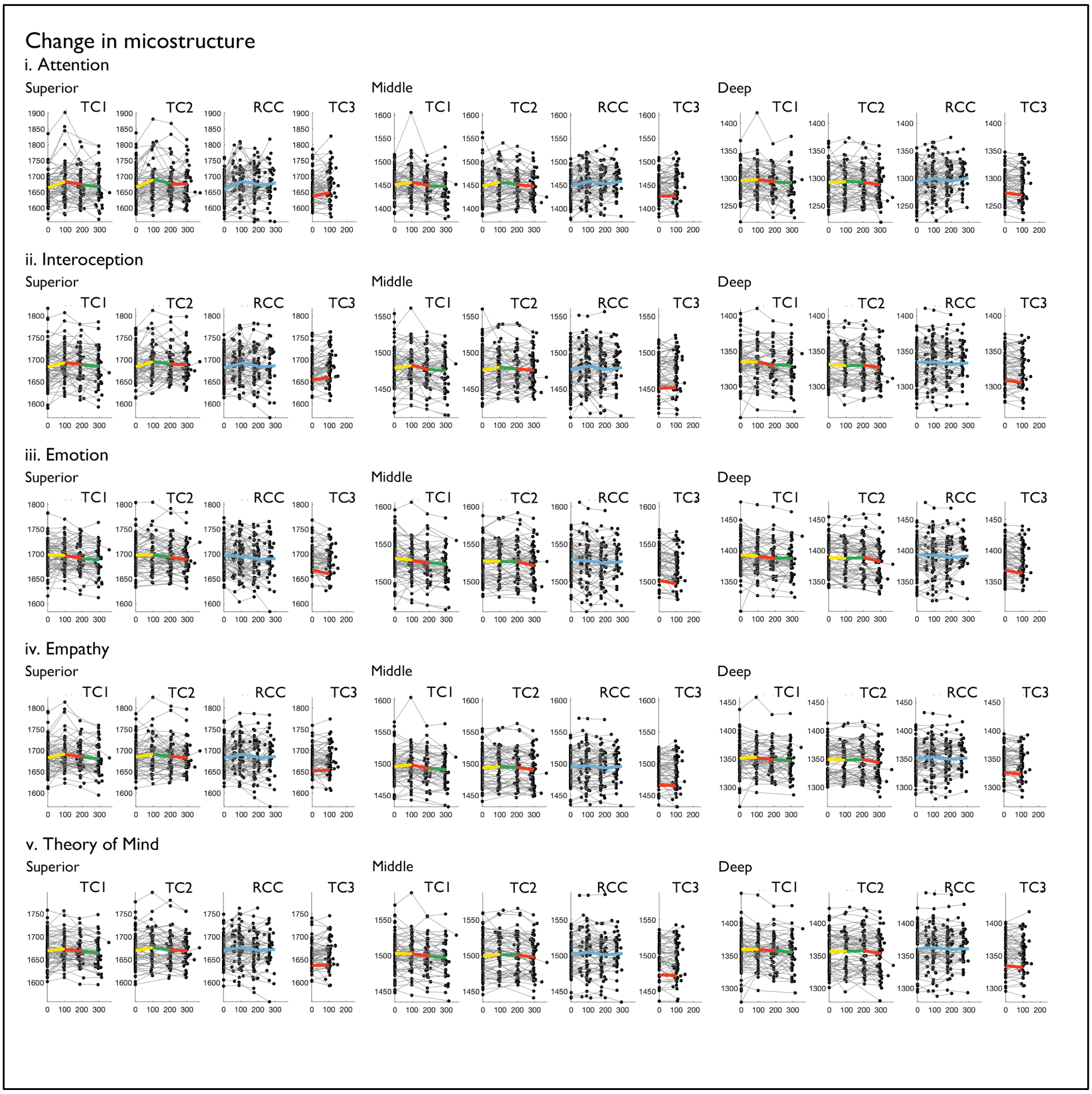

**Supplementary Figure 5. Module-specific change in dept-compartment microstructure as a function of training cohort and functional network.** Yellow = *Presence*, Red = *Affect*, Green = *Perspective*, Blue = *Retest Control*. Changes are displayed as a function of days past baseline (MRI measurement points).

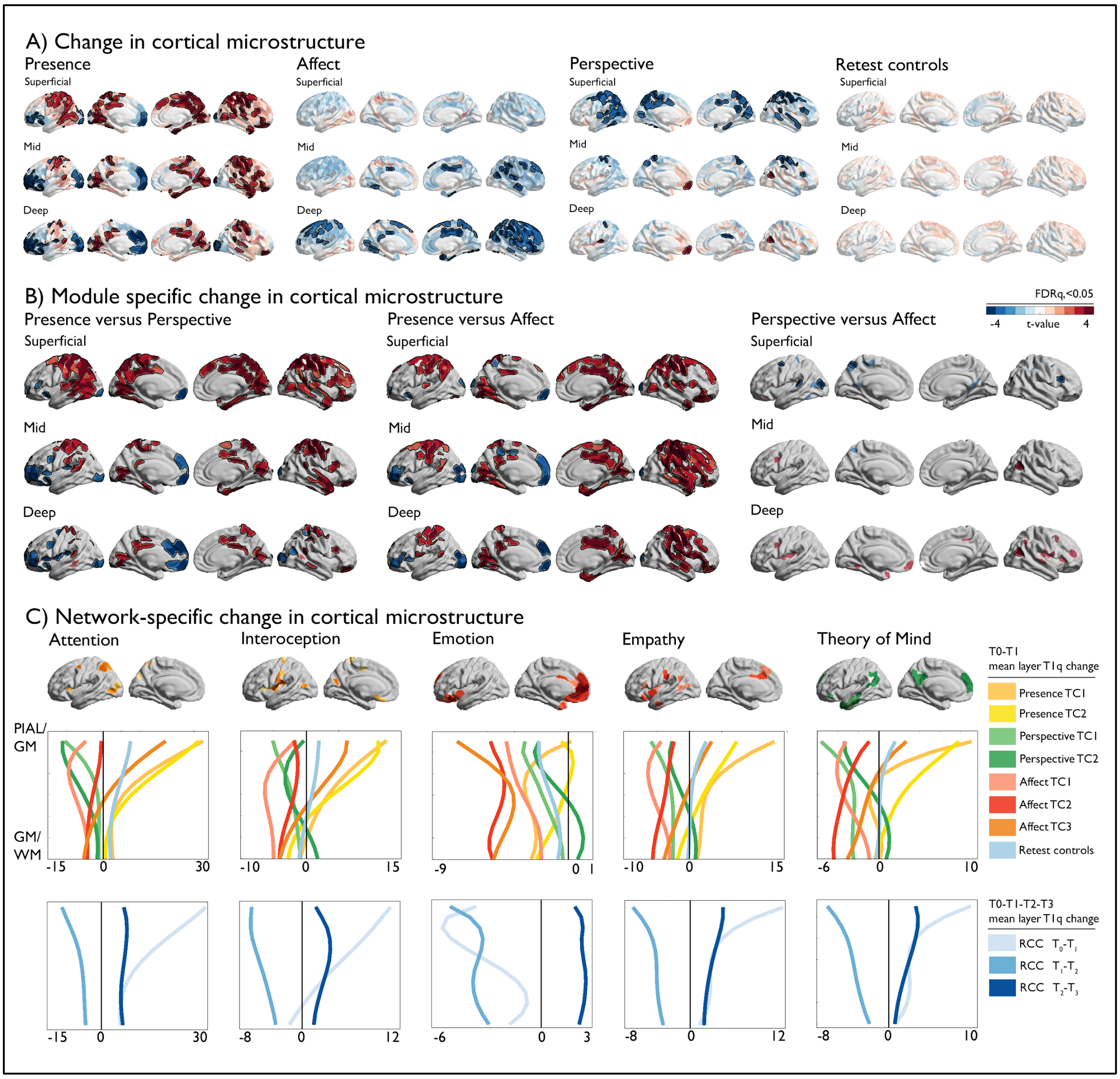

**Supplementary Figure 6. Module-specific change in T1q as a function superficial (1:4) mid (5:8) and deep (8:12) dept compartment.** A). Changes per Module and in retest controls, t-values are projected on the surface and FDRq<0.05 findings outlined in black; B). *Left*: Change in Presence vs Perspective, *middle:* change in Presence versus Affect, *right*: Change in Perspective versus Affect. Trends at p<0.01, FDRq<0.05 outlined in black; C). Network specific changes in microstructure as a function of dept per training cohort, and time point.

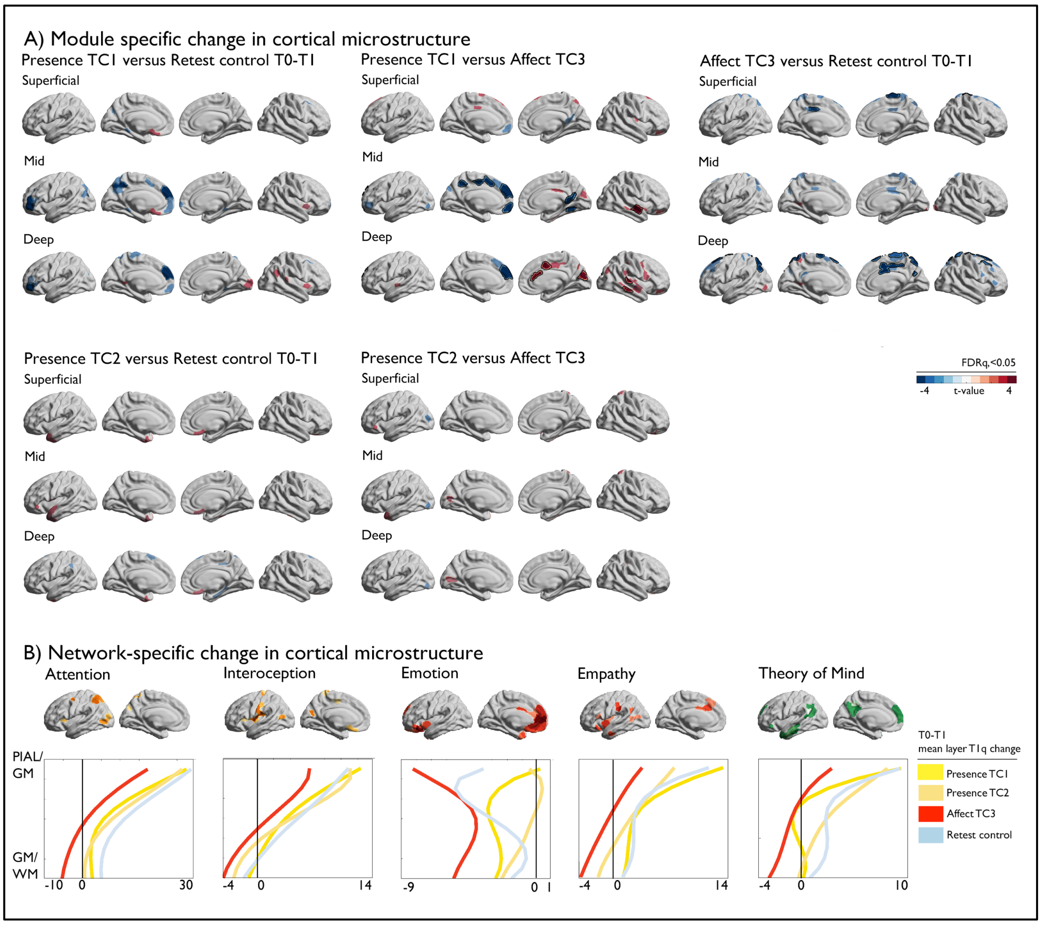

**Supplementary Figure 7. Cohort-specific change in T1q from baseline to T1 as a function superficial (1:4) mid (5:8) and deep (8:12) dept compartment.** A). First row: *Left*: Change in Presence TC1 vs retest controls, *middle:* change in Presence TC1 versus Affect TC3, *right*: Change in Affect TC3 versus retest controls; second row: *Left*: Change in Presence TC2 vs retest controls, *right:* change in Presence TC2 versus Affect TC3, trends at p<0.01, FDRq<0.05 outlined in black; B) Network-specific change in T1q as a function of training cohort in T0-T1.

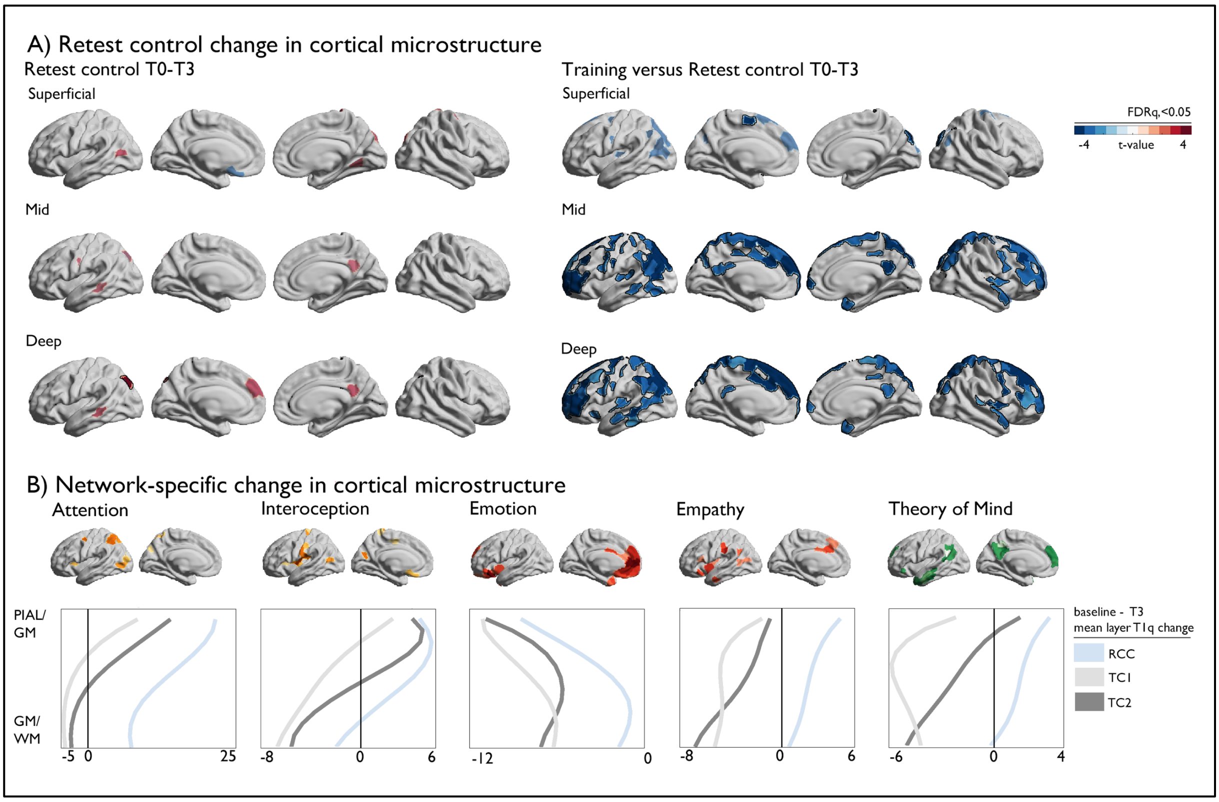

**Supplementary Figure 8. Training-specific change in T1q from baseline to T3 as a function superficial (1:4) mid (5:8) and deep (8:12) dept compartment.** A). *Left*: Change in retest controls, *right*: Change in training versus retest controls, trends at p<0.01, FDRq<0.05 outlined in black. B) Network-specific change in T1q as a function of training cohort.
